# Yuanyang allele pairs enable engineered obligate co-inheritance for population replacement and suppression

**DOI:** 10.1101/2025.11.14.688408

**Authors:** Yunfei Li, Zhuangzhuang Qiao, Chunjie Xian, Shizhe Hu, Jinfeng Jia, Liping Yang, Wuping Li, Miaomiao Tian, Rubing Yang, Qiuyuan Zhang, Hongxia Hua, Jackson Champer, Weihua Ma, Zhihui Zhu

## Abstract

Mendel’s law of segregation, which dictates independent assortment of alleles, is a cornerstone of genetics. Here, we present the Yuanyang allele pair (YYAP), an engineered underdominance gene drive system that enforces obligate co-dependency between homologous alleles via synthetic toxin–antitoxin circuits. This design produces a striking deviation from classical Mendelian segregation: YYAP alleles cannot segregate independently, reshaping inheritance outcomes without altering the physical mechanics of meiosis. In *Drosophila melanogaster*, optimized YYAP strain *K*711 exhibits >99% hemizygous lethality when crossed with wild type, while maintaining stable transmission and high fitness in transheterozygotes. Cage experiments demonstrate efficient population replacement, positioning YYAP as a confined, resistance-proof alternative to CRISPR homing gene drives. Release of only males represents a self-limiting suppression strategy that was also successful in cage experiments. By imposing a near-complete postzygotic incompatibility, YYAP establishes a programmable framework that not only supports pest management, but also enables modeling of reproductive isolation and allelic co-dependency, thereby creating opportunities to explore speciation and broader synthetic inheritance systems in ecology and evolution.

## Introduction

The foundation of classical Mendelian genetics rests upon two fundamental laws: the law of segregation and the law of independent assortment. These principles, first articulated by Gregor Mendel (1866), describe the predictable patterns of inheritance observed during sexual reproduction. In particular, the law of segregation posits that during gamete formation, the two alleles for each gene segregate independently, resulting in each gamete receiving only one allele per gene locus. This process is underpinned by the physical separation of homologous chromosomes during meiosis (Sutton, 1903), ensuring genetic diversity through independent assortment of traits in offspring.

However, the emergence of synthetic biology and advanced genome engineering tools has enabled the design of genetic systems that challenge these long-standing principles. Gene drives represent a class of “selfish” genetic elements engineered to bias their inheritance, spreading rapidly through populations even if they reduce individual fitness. These systems achieve super-Mendelian inheritance (transmission rates >50%) via molecular mechanisms that either enable copying the drive allele or kill offspring lacking the target allele. Endonucleases-based homing drives, such as CRISPR (Clustered Regularly Interspaced Short Palindromic Repeats), are used to cut homologous loci, coercing the cell to copy the drive element onto the damaged chromosome during repair (Hay et al., 2021; Verkuijl et al., 2022; Wang et al., 2022). Toxin-antitoxin systems like *Medea* (Maternal-Effect Dominant Embryonic Arrest) exploit maternal effects: offspring inheriting the drive survive via an antitoxin, while those without it succumb to a toxin (Chen et al., 2007). In CRISPR-based toxin-antidote systems, other variations are possible (Oberhofer et al., 2019; Champer et al., 2020a; Champer et al., 2020b). Additional strategies include X-chromosome shredders that distort sex ratios and transposon-mediated drives (Simoni et al., 2020; Widen et al., 2023). Collectively, these technologies hijack cellular machinery to reshape Mendel’s first law, enabling directed evolution within target populations.

The promise of gene drives lies in their potential to revolutionize pest and vector control. By spreading deleterious traits (e.g., female sterility or pathogen resistance) through wild populations, they could suppress agricultural pests, eliminate invasive species, or disrupt disease transmission by mosquitoes (Yan et al., 2023). Unlike insecticides, which require repeated applications and face widespread resistance, gene drives offer a potentially self-sustaining, species-specific solution with the possibility for permanent population suppression after minimal releases. Nevertheless, critical limitations constrain their real-world applicability. Mutations prevalent in wild populations can compromise the efficacy of both endonuclease-based drives and base-pairing-dependent systems (such as *Medea*). Nuclease-based systems are also sensitive to target-site mutations via end joining (Hay et al., 2021; Verkuijl et al., 2022; Wang et al., 2022). Second, gene drives face ecological and governance challenges. Uncontrolled spread in self-sustaining homing-based systems raises concerns about unintended consequences, such as disrupting food webs, or enabling secondary pests. The self-propagating nature of such global drives also complicates containment, creating transboundary ethical and regulatory dilemmas (James et al., 2023). To circumvent these challenges, modern homing drives leverage optimized CRISPR systems with germline-specific promoters and multiplexed gRNAs to enhance homing efficiency while suppressing resistance allele formation (Champer et al., 2018; Noble et al., 2019; Champer et al., 2020c; Champer et al., 2020d). Split-drive architectures physically separate Cas9 and gRNA components to allow confinement, while daisy-chain drives employ multi-tiered designs for stronger but still self-limiting propagation (Oberhofer et al., 2021).

In 2001, Davis et al. theorized an elegant underdominance (UD) implementation using two interdependent genetic constructs, each carrying a toxin and the antitoxin for the other toxin. When located at the same genomic locus on homologous chromosomes, these constructs would render hemizygotes nonviable (due to lacking one antitoxin), while transheterozygotes would persist. This design explicitly required the simultaneous inheritance of both alleles located on the homologous chromosome pair, creating functional inseparability at the locus level. This and similar systems have been the subject of extensive computational modeling due to their desirable features (Magori and Gould 2006; Huang et al., 2009; Huang et al., 2011; Edgington and Alphey 2017; Dhole et al 2018; Champer et al., 2020e; Sánchez et al 2020). Despite their conceptual promise, such two-construct systems have never been experimentally realized in any organism due to the extreme difficulty of engineering tightly linked, interdependent genetic modules without disrupting essential functions. Similar designs have been developed, but some involve engineered *Medea* elements that function through maternal-effect toxicity in heterozygous females (where hemizygous males are viable), and do not function outside *Drosophila*, or remain at the design/modeling stage (Akbari et al., 2013; Champer et al., 2020e; Li and Champer 2023).

Here, we transcend these limitations by creating the first biological system where homologous alleles are biologically inseparable: the Yuanyang Allele Pair (YYAP; Yuanyang refers to the Mandarin duck *Aix galericulata*, a cultural symbol of inseparable partnership in Chinese culture) in *Drosophila melanogaster* (Figure 1). This design ensures that allelic components P and M on identical loci of homologous chromosomes are inherited as a unit. We engineered the YYAP platform utilizing toxin-antitoxin (TA) systems, which are genomically ubiquitous in bacteria and functionally active across diverse eukaryotes including insects (de la Cueva-Méndez et al., 2003; Slanchev et al., 2005; Zhang et al., 2021; Li et al., 2025). The top-performing YYAP carrier exhibited stable inheritance, and when crossed with wild-type (WT) strains, >99% of the offspring died by the third instar. Cage tests showed that YYAP enables efficient gene drive-based population replacement. Repeated releases of males can also support a self-limiting population suppression strategy. The YYAP system represents the first experimental realization of mandatory co-inheritance of homologous alleles, challenging Mendel’s law of segregation and offering unprecedented advantages in pest control by eliminating drive resistance, avoiding ecological risks, and enabling dual control strategies.

**Figure 1.**
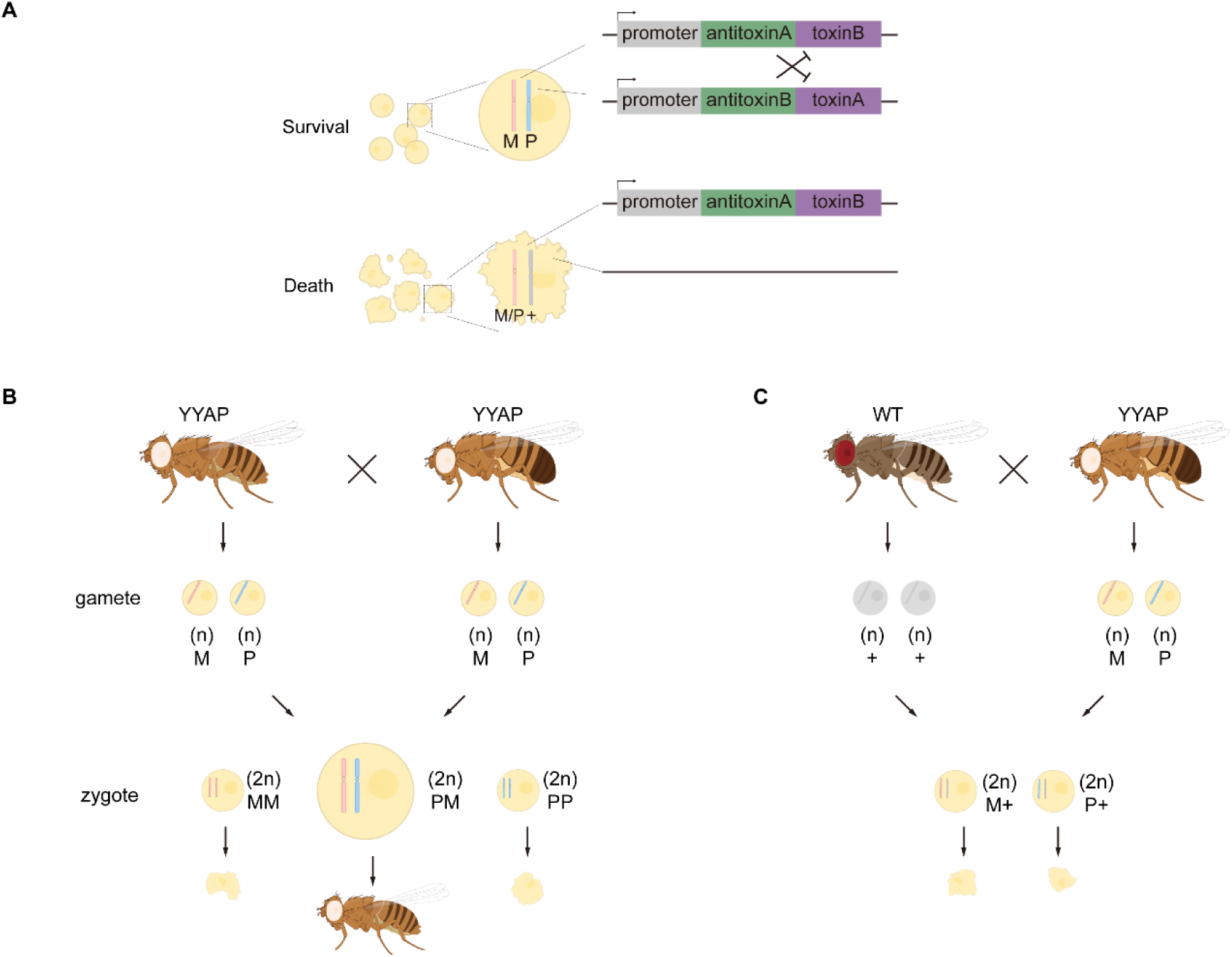
Overall design and implementation of the Yuanyang Allele Pair (YYAP) system. (A) Schematic representation of the YYAP principle. YYAP consists of the paternal P allele and the maternal M allele, constructed on a toxin–antitoxin system. The P allele carries antitoxin B-toxin A, while the M allele carries antitoxin B-toxin A. When both alleles are present, toxins are neutralized by the corresponding antitoxins, allowing cell survival; when separated, one allele expresses a toxin gene while the wild-type (WT) homolog lacks the corresponding antitoxin, leading to cell death. (B and C) Expected behavior of YYAP strains (*yellow*) during self-crossing and mating with wild-type flies. Gray fly denotes WT, damaged cells denote non-viable zygotes. P, allele expressing antitoxin B-toxin A; M, allele expressing antitoxin B-toxin A; +, WT allele; n, haploid gamete; 2n, diploid zygote.

## Results

### Construction and Optimization of YYAP Transgenic strains in Drosophila

We developed the YYAP system using TA modules (Figure 1). In our previous work, we identified several type II TA pairs in Sf9 and S2 cell lines that exhibited effective toxicity toward insect cells, with cognate antitoxins effectively neutralizing the corresponding toxins. Notably, no cross-neutralization was observed among different pairs (Zhang et al., 2021; Li et al., 2025). Initial proof-of-principle experiments in Sf9 cells confirmed the design: co-transfection mimicking YYAP state showed normal viability, whereas simulated allelic separation caused near-complete cell death (Figures S1, and 1A). To implement YYAP in *D. melanogaster*, we engineered interdependent genetic modules (antitoxin B–toxin A and antitoxin A–toxin B) precisely inserted at homologous loci (86Fb) (Figure 1A, Table S1). To suppress unintended toxin expression during strain construction, we placed a removable selection marker cassette—flanked by FRT sites—between the TA coding sequence and its promoter (Figure S2A). Distinct FRT variants (FRT3 or FRT5) were used on homologous chromosomes to avoid ectopic recombination. Using φC31-mediated site-specific integration, we established two parental strains: strain *P’* (antitoxin B–toxin A) and strain *M’* (antitoxin A–toxin B). Crossing these strains, followed by heat-shock–induced FLP recombinase activation, triggered excision of the selection cassette and restored TA expression, yielding YYAP chimeric flies (Figures 2A, S3B and S3C). Germline-recombined progeny were subsequently screened to establish stable YYAP strains (Figure S3D).

**Figure 2.**
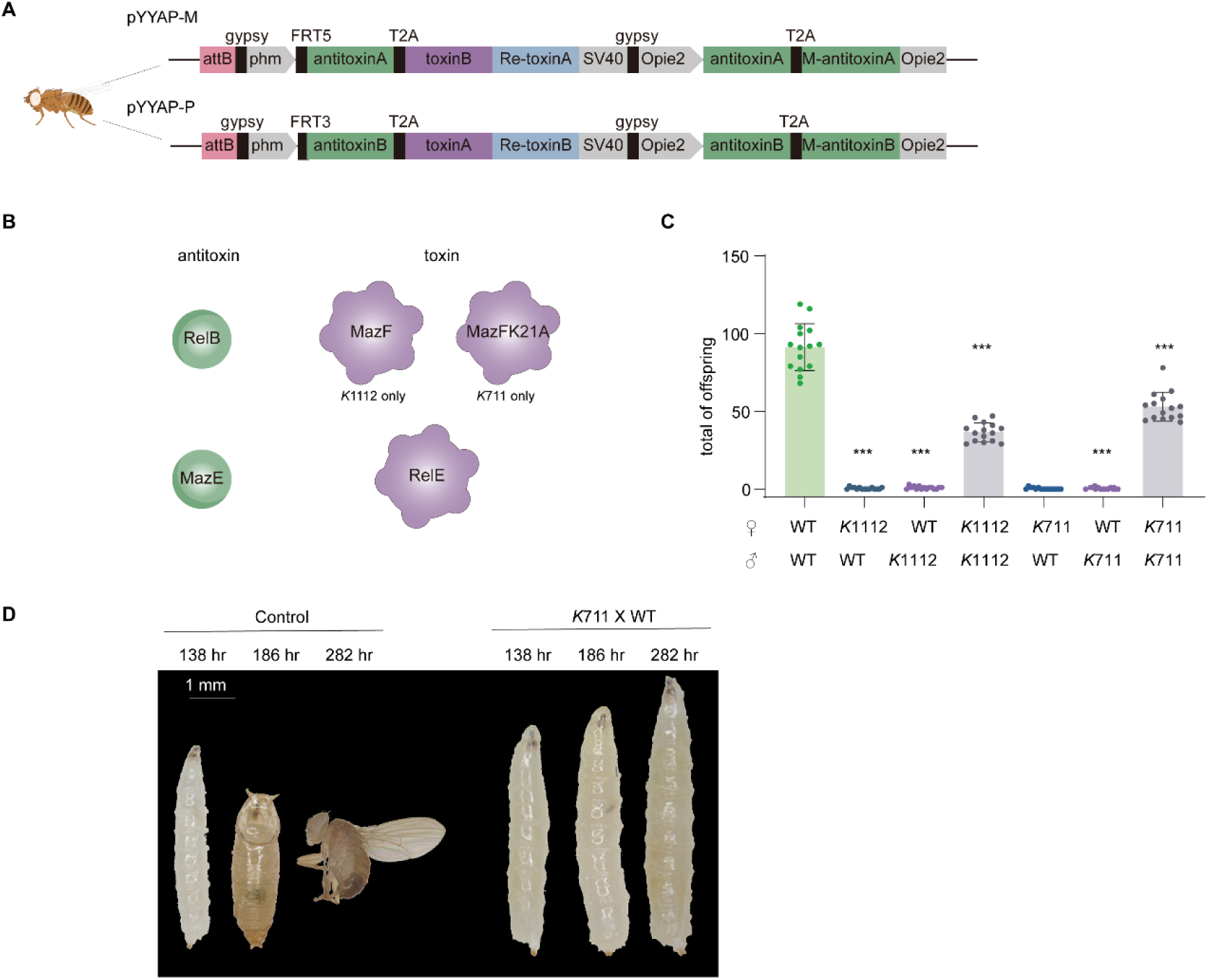
Lethal effects of the YYAP system. (A) Schematic of the transgenic insertion element in *Drosophila* based on the type VIII transgenic construct used in this study. The attB site mediates site-specific integration via φC31 recombinase; the *phm* promoter drives expression of the antitoxin-T2A-toxin cassette; Acie1::antitoxin-Op64 provides homologous antitoxin genes to counteract leaky toxin expression; *yellow/miniwhite* markers facilitate line identification; FRT3/5 sites enable FLP recombinase-induced DNA excision of markers and antitoxin cassettes; the Antitoxin-T2A-toxin cassette uses a 2A peptide for bicistronic translation; Re-toxin generates antisense RNA; Opie2::antitoxin-T2A-antitoxin-Opie2 ensures constitutive antitoxin expression; the gypsy insulator blocks potential cis-regulatory activity near the cassette. (B) Schematic representation of the toxin–antitoxin modules, with the green module indicating the antitoxin and the purple module indicating the toxin. (C) Hemizygous lethality assay showing the number of progeny surviving to adulthood. Genotypes of parental strains are indicated on the x-axis. Statistical comparisons were performed using Welch’s *t*-test (unequal variances) against the control group, *n* = 15. (D) Phenotype of WT (left) and hemizygous offspring from *K*711 × WT (right) at three developmental time points. Scale bars, 1 mm. Statistical significance: * *p* < 0.05, ** *p* < 0.01, *** *p* < 0.001.

Given the complexity of TA expression regulation in eukaryotic systems, we performed iterative vector optimization from design Types I to VIII (Figure S2). Key improvements included: Type I established the core antitoxin-toxin cassettes, but suffered from leaky expression; Type II added Acie1(promoter)-driven antitoxins between FRTs to neutralize toxin leaky expression from transcription readthrough; Type III incorporated microRNA to reduce toxin mRNA level; Type IV and VI increased antitoxin copy numbers to boost expression; Type V and VI changed promoter for antitoxin expression; Type VII uncoupled the expression of antitoxin with toxin; Finally, Type VIII introduced inverted toxin sequences to generate antisense RNA and reduce toxin transcript levels, mutated MazF to attenuate toxicity (for strains *K*711 and *K*811), and combined all other optimizations, achieving near-zero background toxicity. We also evaluated multiple promoters, including *serendipity α* (*sry-α*) and *phantom* (*phm*) etc., across diverse TA combinations. In total, we constructed 76 plasmids and performed embryo injections (Table S2). Transgenic strains were validated by both phenotypic screening of the marker and genotypic analysis confirming the presence of the exogenous gene and its insertion into the 86Fb site (Figure S4A; additional validation data available upon request). Of these, 44 strains (22 pairs) met the criteria for further analysis (Table S3). However, during mating and heat-shock–induced recombination (Figure S3), many strains failed due to infertility, reduced progeny fitness, or absence of mosaic individuals (Figure S5). Ultimately, only nine strains completed the entire pipeline, yielding stable YYAP *Drosophila* strains with robust expression and reliable inheritance (Table S3, also see following section).

### Identification of YYAP Strains with Strong Hemizygous Lethality and Optimal Transheterozygous Fitness

The core requirement of YYAP is to enforce non-segregation of homologous alleles, which predicts two outcomes: (i) self-crosses of YYAP strains yield exclusively transheterozygotes, with a theoretical survival rate of ∼50%; and (ii) crosses with wild-type (*y*^1^*w^1^*, unless otherwise noted) result in complete hemizygous lethality due to unopposed toxin exposure. Genetic evaluation of the nine YYAP strains revealed marked variation in performance. Seven strains (*S*914, *S*1718, *D*1112, *K*1718, *K*914, *K*67, and *K*811) produced homozygous offspring (Figures S5A-S5C), indicating insufficient toxin potency or expression, thereby compromising the YYAP mechanism. Consistently, these strains exhibited weak or absent lethal effects when crossed with WT (Figures S6A–S6F, and S6M).

In contrast, strains *K*1112 and *K*711 demonstrated robust hemizygous lethality. In reciprocal crosses with WT, >99% of progeny died at the third instar larval (L3) stage, with only a small fraction of surviving adults (0.36-0.95%) (Figures 2C and S6I-S6J; Table S4). Genotyping confirmed that survivors were hemizygotes without detectable mutations in toxin sequences or regulatory regions: in the *K*1112 group, all surviving individuals were of the *M* [MazE2783 (*Escherichia coli*)–RelE1223 (*Streptococcus pneumonia*)]*/+* genotype, whereas in the *K*711 group, survivors included both *M* [MazE2783 (*E. coli*)–RelE1223 (*S. pneumonia*)]*/+* and *P* [RelB1224 (*S. pneumonia*)–MazF2782K21A (*E. coli*)]*/+* hemizygotes carrying the WT allele (Figure S4C-S4D). Hemizygous survivors crossed again with WT produced ∼50% viable offspring (Figure S6G). Reconstructed *K*711* strains from these surviving hemizygotes, followed through up to ten generations of self-crossing, consistently maintained high hemizygous lethality (Figure S6H), excluding epigenetic silencing as the predominant escape mechanism. The low survival frequency likely reflects stochastic variation in toxin promoter activity or individual differences in sensitivity.

Despite its strong hemizygous lethality, *K*1112 displayed severe fitness defects. Eclosion rates were significantly reduced compared to WT (53.23 ±3.15% vs. 86.64 ±1.50%, *p* < 0.001; Figure S6K), self-cross offspring viability was well below theoretical expectations (40.07% vs. expected 50%, *p* < 0.001; Figure 2C), and sex ratios were skewed (31.12% males vs. expected 50%, *p* < 0.001; Figure S6L). These fitness defects might result from insufficient toxicity neutralization and sex imbalance likely reflects sex-specific differences in *phm* promoter copy number in *K*1112 (four copies in females vs. three in males), leading to disproportionate toxin expression. By contrast, *K*711, constructed with the attenuated *E. coli* toxin MazFK21A, exhibited optimal performance. It maintained >99% hemizygous lethality (Figure 2C), while overall fitness was indistinguishable from WT: eclosion rates (88.69 ±0.87%) were comparable to WT (86.64 ±1.50%, *p* = 0.585), self-cross offspring viability exceeded theoretical expectations (57.95% vs. expected 50%, *p* < 0.001; Figure 2C), likely due to enhanced resource availability, and sex ratios were balanced (*p* = 1.000; Figure S6L). Importantly, YYAP strains demonstrated genetic stability across generations, as exemplified by *K*711: RT-PCR confirmed normal transcription of toxin–antitoxin cassettes (Figure S4B), and both genotyping (*n* = 46 at G5, *n* = 90 at G10) and sequencing (*n* = 3 at both G5 and G10) revealed 100% *P/M* transheterozygotes without mutations. Collectively, these results establish *K*711 as the ideal YYAP strain for subsequent functional studies.

### Hemizygous Lethality and Biological Features of the K711 strain

The *K*711 strain employs the *phm* promoter, which is specific to the prothoracic gland (PG), to drive expression of the nucleases MazFK21A (*E. coli*) and RelE (*S. pneumoniae*) (Warren et al., 2002; Petryk et al., 2003). The PG is essential for producing the molting hormone 20-hydroxyecdysone (20E). Hemizygous hybrids (*P/+* or *M/+*) from *K*711 × WT crosses consistently arrested at L3 stage, with only 0.36–0.95% progressing to pupae or adults. Arrested larvae exhibited continuous growth and significantly enlarged body size (Figure 2D), a hallmark of ecdysone deficiency (Colombani et al., 2005; Yamanaka et al., 2015; Liu et al., 2024). Developmental timing analyses confirmed that lethality was confined to the L3 stage (Table S4).

Fecundity and hatchability varied depending on cross direction. On the first three days, *K*711 females ×WT males laid 151.8 ±7.1, 635.1 ±34.8, and 760 ±30.7 eggs, respectively, with a hatchability of 79.4 ±2.0%. In contrast, *K*711 males ×WT females produced 268.7 ± 17.6, 771.9 ± 47.6, and 1028.4 ± 34.8 eggs on the first three days, respectively, but had lower hatchability (61.6 ±3.3%) (Figures 3A and S7A). Despite these differences, L1 larval numbers remained ∼79% of controls. Across genotypes (*+/+*, *P/+*, *M/+*, *P/M*), no differences were observed in L1–L2 duration, and *P/M* larvae showed L3 timing comparable to *+/+*. As expected, nearly all *P/+* and *M/+* hemizygotes arrested at L3 (Table S4). Measurements of larval size at 138, 186, and 282 hours post-oviposition revealed that *P/+* and *M/+* larvae grew significantly larger than WT at early time points, and later exhibited divergent trajectories: either continued feeding and growth, or reduced activity with stable weight and length (Figures S7B and S7C). *P/M* pupae displayed prolonged pupal duration, likely reflecting residual toxin activity delaying ecdysone accumulation (Table S4).

**Figure 3.**
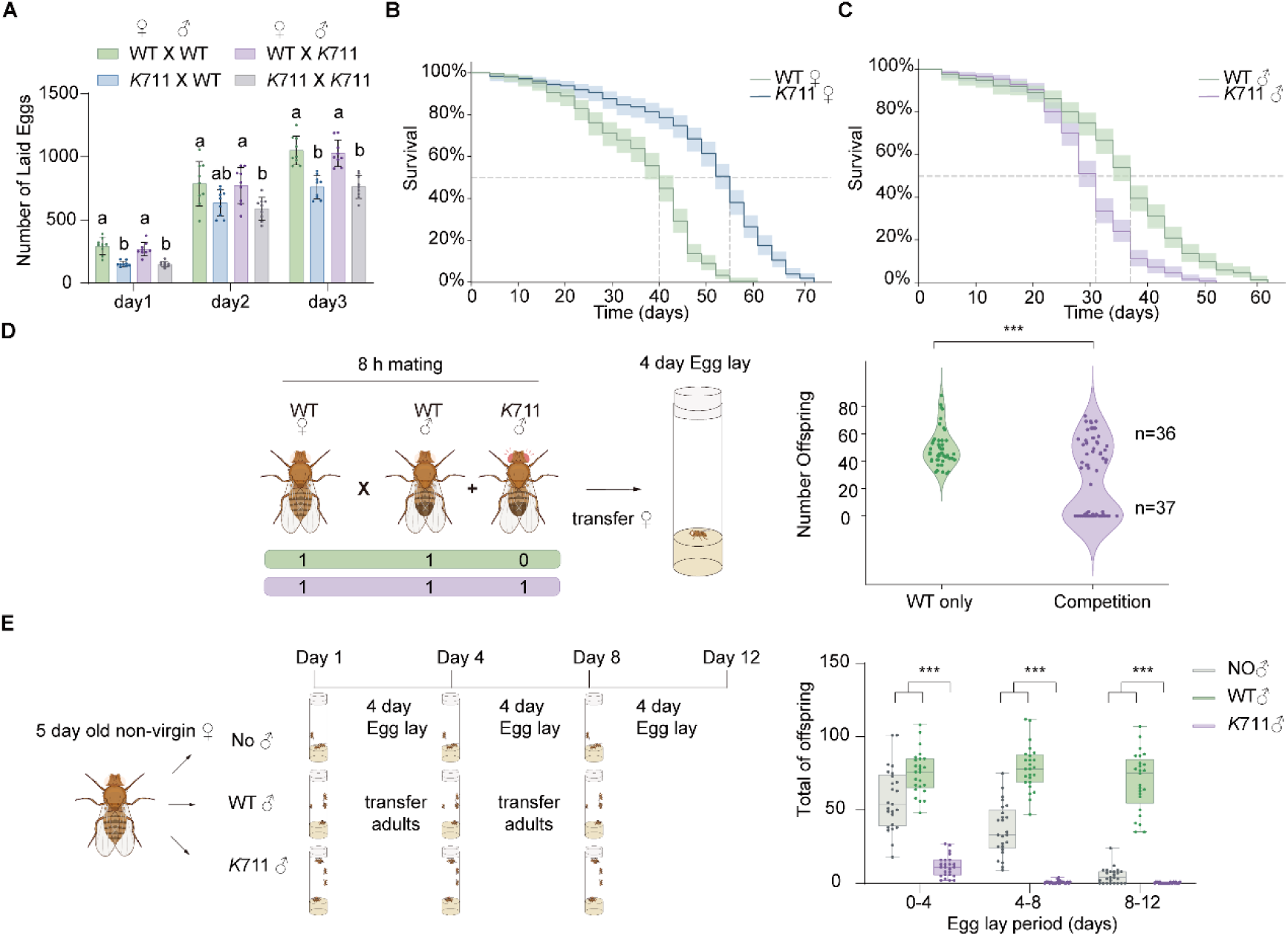
Fitness measurements of *K*711 YYAP. (A) Egg production of females within 24 hr over three consecutive days. A one-way ANOVA followed by Tukey’s multiple comparisons was performed on the egg production of females across four mating groups (*n* = 9). (B) Survival curves of WT females (*N*=3, *n*=243, green line) versus *K*711 females (*N*=3, *n*=257, blue line). (C) Survival curves of WT males (*N*=3, *n*=258, green line) versus *K*711 males (*N*=3, *n*=262, purple line). (D) Experimental design (left panel) and results (right panel) of individual male mating competition assay. Violin plots show the number of offspring produced by control group (WT♀ × WT♂, green, *n* = 45) and the competition group (WT♀ × WT♂ + *K*711♂, purple, *n* = 74). Statistical comparisons were performed using Welch’s t-test (unequal variances). (E) Experimental design (left panel) and results (right panel) of sperm displacement assay. Box plots represent non-virgin WT females without additional males (gray), with WT males (green), or with *K*711 males (purple). For each egg-laying period, Welch’s *t*-tests were performed to compare the *K*711 group with both the no-male and WT male groups (*n* = 25 per group). Survival data are presented as Kaplan–Meier probability estimates with 95% Greenwood confidence intervals (shaded areas). The dashed line indicates the median survival time at 50%. Group differences were determined using the log-rank test. Statistical significance: * *p* < 0.05, ** *p* < 0.01, *** *p* < 0.001. Letters in panels indicate significance at α = 0.05.

Because YYAP’s application in pest control depends on adult reproductive performance, we also assessed lifespan. WT median lifespan were 40 days for females (95% CI: 37–43, *N* = 3, *n* = 243) and 37 days for males (95% CI: 34–37, *N* = 3, *n* = 258) (Figures 3B and 3C). *K*711 females exhibited significantly longer lifespan (median 55 days, 95% CI: 52–55, *N* = 3, *n* = 257, *χ²* = 183.36, *p* < 0.005) (Figure 3B), whereas *K*711 males had reduced lifespan (median 31 days, 95% CI: 28–31, *N* = 3, *n* = 262, *χ²* = 68.55, *p* < 0.005) (Figure 3C). Although WT lifespans vary with diet and temperature (Lin et al., 1998; Clancy et al., 2001; Tatar et al., 2001), the cause of extended female longevity in *K*711 remains unclear. In contrast, reduced male lifespan is likely attributable to residual toxin activity insufficiently neutralized by the antitoxin.

### Mating Competitiveness and Sperm Displacement by K711 Males

The effectiveness of population suppression strategies depends on YYAP males competing successfully with WT males for mates. We assessed the competitiveness of *K*711 males using two established assays (Kandul et al., 2019; Upadhyay et al., 2022). In single-pair competition assays (1 *K*711 male + 1 WT male + 1 WT female, 8 hours), offspring counts over the first four days displayed a bimodal distribution: thirty-six females produced numerous WT adults (indicating successful WT mating), while thirty-seven females produced few or no viable adults (indicating successful *K*711 mating, with progeny dying from hemizygous lethality during L3) (Figure 3D). Observed L3 larvae from the thirteen females exhibited canonical YYAP lethal phenotypes, demonstrating that *K*711 males compete as effectively as WT males. In group mating assays, 20 WT females were housed with varying male ratios (0:4, 1:3, 2:2, 3:1; *K*711:WT). As the proportion of *K*711 males increased, viable adult offspring declined significantly (Figure S8A), confirming effective competitiveness of *K*711 males under group mating conditions.

Because *Drosophila* females can store sperm from multiple matings and engage in sperm competition (Brown et al., 2004; Castrezana et al., 2017), we next tested the sperm displacement ability of *K*711 males (Clark et al., 1995; Upadhyay et al., 2022). In continuous displacement assays, non-virgin females previously mated with WT males were subsequently co-housed with *K*711 males. Compared with controls (no males or only WT males), *K*711 exposure reduced viable offspring by 80.19%, 93.60%, and 97.71% across three oviposition periods (Figure 3E), indicating that *K*711 sperm effectively displaced or suppressed stored WT sperm. In time-limited competition assays, *K*711 males functioned as either attackers (displacing existing WT sperm) or defenders (resisting WT displacement). In both contexts, viable offspring were significantly reduced—by 57.47% and 77.19%, respectively (Figure S8B)—highlighting the strong competitive efficiency of *K*711 sperm.

Together, these results establish that *K*711 males exhibit mating competitiveness comparable to WT males, while displaying superior sperm competition and displacement capacity. These behavioral attributes provide a solid foundation for the deployment of YYAP in population control strategies.

### Cage Experiments Demonstrate K711-Mediated Population Replacement and Suppression

We next evaluated the practical potential of YYAP in controlled cage populations (*Drosophila*), following the experimental design outlined in Figures S9A and S10. For population replacement, we released both *K*711 males and females (1:1 sex ratio) at varying initial WT:*K*711 ratios (Figures 4A and S9). At low release ratios (WT:*K*711 = 1:1), *K*711 frequencies declined rapidly and were fully replaced by WT within 3–7 generations (Figures 4A and S9D). This outcome reflects the inherent reproductive disadvantages of YYAP individuals: self-crosses yield 50% lethal progeny, while crosses with WT result in complete lethality. At medium ratios (*K*711:WT = 5:1), two of three replicates achieved full replacement by generation 6–7, whereas the third went extinct, underscoring the sensitivity of replacement outcomes to initial releasing ratios (Figures 4A and S9E). High-ratio releases (*K*711:WT = 10:1) consistently drove rapid and stable replacement, with *K*711 frequencies reaching 100% by generation 4 and maintained thereafter (Figures 4A and S9F). Post-replacement populations stabilized at ∼50% of their original size (Figure S9F), this reduced value because half of offspring would be nonviable.

**Figure 4.**
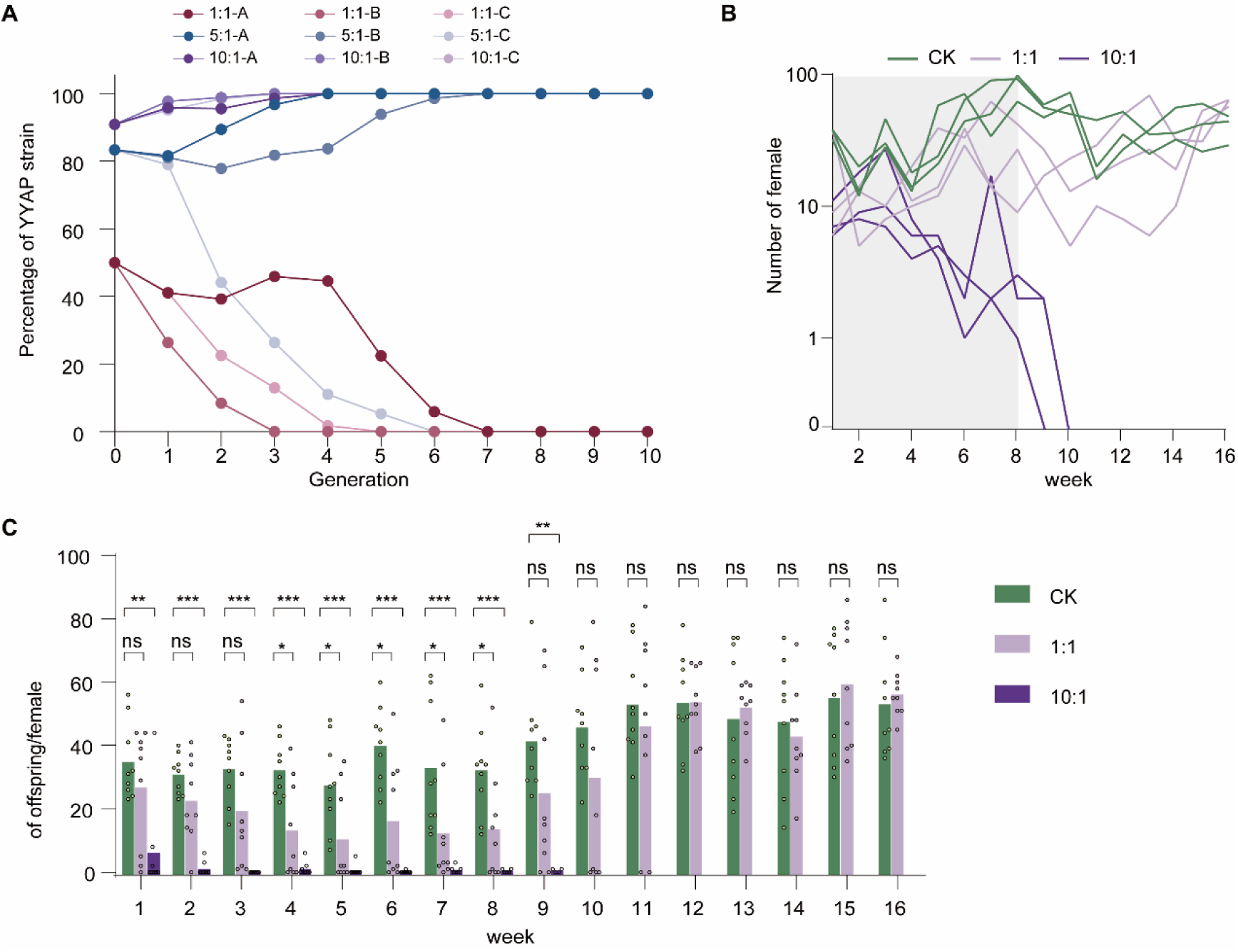
Laboratory cage trials of population suppression and replacement of wild-type *D. melanogaster*. (A) Dynamics of population replacement in small, non-overlapping generation cages at different *K*711 release ratios. Small cages were seeded with 15 WT females and 15 WT males, along with additional *K*711 females and males to achieve release ratios of 1:1, 5:1, or 10:1 (15♀ + 15♂, 75 ♀ + 75♂, or 150♀ + 150♂ *K*711 flies, respectively). Offspring in subsequent generations were scored as RFP-positive (due to a 3×P3::RFP cassette upstream of the insertion site) or WT. The y-axis represents the percentage of RFP flies in the total population, and the x-axis represents each generation. Panels (A), (B), and (C) correspond to the 1:1 replacement, 5:1 replacement, and 10:1 replacement groups, respectively. (B) Female population dynamics measured from trapping devices. Light gray shading indicates weekly additional *K*711 releases through week 8. In control cages (green lines), the population increased gradually and plateaued by week 8. In 1:1 suppression cages (light purple lines), populations increased slowly until week 6, then began to decline; in one cage, recovery began at week 10, while in the other two cages, populations started to rise again by week 12. In 10:1 suppression cages (dark purple lines), the number of wild-type females rapidly declined until they became undetectable by week 10. (C) Weekly fecundity assay results. Each colored dot represents the total number of offspring from a single vial containing one female, with colors indicating control cages (green), 1:1 (*K*711:WT) suppression cages (light purple), and 10:1 (*K*711:WT) suppression cages (dark purple). Statistical comparisons between the control group and the different *K*711 release ratio treatments were performed weekly using the Mann–Whitney test (*n* = 2–9 per group). Statistical significance: ns, not significant; * *p* < 0.05, ** *p* < 0.01, *** *p* < 0.001

Importantly, unlike population replacement strategies based on cytoplasmic incompatibility (Zheng et al., 2019), inadvertent release of YYAP females does not compromise suppression: their crosses with WT males also produce non-viable offspring. Moreover, at low release ratios, female introductions do not promote population replacement (Figure 4A). Together, these results demonstrate that high-ratio *K*711 releases can achieve both effective population suppression and rapid, stable population replacement, supporting YYAP as a versatile genetic control strategy.

For population suppression (*K*711 male releases only), we tested two release strategies. In low-ratio releases (100 *K*711 males weekly; *K*711:WT = 1:1), suppression was transient and populations rebounded following releases (Figures 4B, S10B and S10C), indicating insufficient eradication pressure. By contrast, high-ratio “overflooding” releases (1000 *K*711 males weekly; *K*711:WT = 10:1) produced rapid suppression: female fertility dropped significantly within the first week (Figure 4C), and despite transient population increases due to mass releases, the strong hemizygous lethality of *K*711 led to a swift collapse, with complete extinction achieved by week 14 (no adults detected; Figures 4B, S10B and S10C).

### Modeling Predicts YYAP as a Threshold-Dependent Strategy for Mosquito

#### Population Control

Given the activity conservation of TA in eukaryotes and the demonstrated efficacy of YYAP in *Drosophila*, we applied the MGDrivE framework to simulate YYAP deployment in mosquito populations (*Aedes aegypti*) (Sánchez et al., 2019). Simulations modeled a population of 10,000 adult females over one year, incorporating weekly YYAP male-only releases for six months to evaluate suppression, and single or double releases of both sexes to assess population replacement (parameters in Table S5). For population suppression, high-ratio male releases (YYAP:WT=10:1) led to complete collapse of wild female populations within 4–6 months (Figure S11A). For population replacement, a single release at a ratio of 1:6.6 (WT:YYAP) and a double release at 1:3.1 achieved fixation, stabilizing populations at ∼50% of baseline levels, consistent with the intrinsic viability threshold of YYAP (Figures S11D and S11E). Sensitivity analyses demonstrated the robustness of these outcomes to reductions in YYAP male lifespan and mating competitiveness (Figures S11B, S11C, S11F and S11G), underscoring the threshold-dependent nature of YYAP dynamics. Collectively, these modeling results support YYAP as a potent, scalable, and threshold-dependent genetic control tool with broad applicability to mosquito-borne disease suppression.

## Discussion

The YYAP system establishes the first experimental paradigm of obligate co-inheritance of homologous alleles, directly challenging the phenotypic outcomes dictated by Mendel’s law of segregation. By engineering synthetic toxin-antitoxin circuits that enforce an inseparable partnership between alleles on homologous chromosomes, we have created a genetic system where inheritance is no longer a probabilistic assortment but a deterministic outcome of allelic co-dependency. In *D. melanogaster*, our optimized YYAP strain K711 achieved the critical design objective: it imposed >99% lethality in hemizygous offspring from crosses with wild-type individuals, while maintaining stable transmission and near-wild-type fitness in transheterozygotes. This work thus transcends prior theoretical proposals and modular modeling of underdominance systems by delivering a functional, resistance-proof, and ecologically confined platform for rewriting inheritance logic, with direct applications in population management and evolutionary modeling.

### Design Principles and Iterative Optimization: From Concept to Biological Reality

The realization of YYAP hinged on solving two fundamental challenges: the precise integration of interdependent genetic modules at identical loci on homologous chromosomes, and the prevention of lethal toxin expression during strain construction. Our solution leveraged a combination of site-specific recombination systems (ΦC31 for precise integration and FLP/FRT for conditional activation), a strategy inspired by synthetic gene circuit design in eukaryotes (Roquet et al., 2016). The journey from our initial Type I vectors to the optimized Type VIII design underscores the intricate balance required for such a system to function in a complex eukaryotic organism. Early constructs suffered from leaky toxin expression, leading to developmental defects. The incorporation of FRT-flanked cognate antitoxin cassettes was a pivotal step, allowing us to suppress toxicity until the final strain was established. Subsequent refinements—including the addition of gypsy insulators, increased antitoxin copy numbers, antisense RNA constructs to degrade toxin transcripts, and the strategic attenuation of toxin activity (e.g., MazFK21A)—were instrumental in achieving strains like *K*711 with near-zero background toxicity (Figure S2; Tables S2 and S3). This systematic optimization, screening 76 constructs across various promoters and TA pairs, epitomizes the “design-build-test-learn” cycle central to synthetic biology and highlights the critical parameters for success: the use of a mitosis-specific promoter (e.g., *phm*) to avoid gametogenesis toxicity, moderate toxin strength to allow for complete neutralization in heterozygotes, and matched potency between the two toxins to prevent homozygous formation (Figure S5; Tables S2 and S3).

### Biological Characterization of *K*711: A Balanced and Robust System

The *K*711 strain, combining the attenuated *phm*-*MazFK21A* and phm-*RelE* cassettes, emerged as our top-performing candidate by striking an optimal balance between stringent hemizygous lethality and minimal fitness cost in carriers (Figures 2C, S6J, S6K and S6L). The characteristic L3 larval arrest phenotype of hemizygous offspring, marked by enlarged size and failed pupariation (Figures 2D; Table S4), is a direct consequence of ecdysone deficiency resulting from the ablation of prothoracic gland function by the toxins (Warren et al., 2004; Liu et al., 2024), confirming the tissue-specific efficacy of our design. The rarity of escapers (<1%), which showed no mutations in toxin sequences or promoters upon genotyping, points to stochastic variations in expression or sensitivity rather than genetic resistance—a finding consistent with bet-hedging strategies in biological systems (Slatkin, 1974; Dobrzyński et al., 2011). The stability of reconstructed *K*711* lines over ten generations, maintaining full lethality (Figure S6H), is particularly noteworthy when contrasted with the propensity for resistance emergence in CRISPR-based homing gene drives (Hammond et al., 2021). While minor fitness costs were observed—such as reduced fecundity in *K*711 females crossed with WT males (Figures 3A) and a modest reduction in male lifespan (Figure 3C)—these are attributable to incomplete toxin neutralization in specific contexts and are significantly less severe than the fitness penalties associated with radiation sterilization used in traditional SIT (Holbrook and Fujimoto, 1970; Suckling et al., 2019). Crucially, *K*711 males demonstrated mating competitiveness and sperm displacement capability indistinguishable from their wild-type counterparts (Figures 3D, 3E, S8A and S8B), a key attribute for any field-type counterparts (Figures 3D, 3E, S8A and S8B), a key attribute for any field.

### Dual-Mode Population Control Management: Confined Replacement as a Primary Advantage

The biological characterization of the *K*711 strain—exhibiting nearly wild-type fitness such as robust mating competitiveness—provides a solid foundation for its practical deployment. Building directly on these attributes, our cage experiments demonstrate that the YYAP system can be strategically configured to achieve two distinct, yet complementary, population management objectives: suppression and replacement.

In suppression mode, YYAP operates as a genetic analogue of the Sterile Insect Technique (SIT), wherein repeated releases of engineered males reduce target population density through reproductive incompatibility. Our data show that a 10:1 release ratio drives complete population collapse within 13–14 weeks—a performance broadly comparable to other advanced genetic systems. For instance, the fsRIDL strain OX3604C achieved elimination in 10–20 weeks at similar ratios (Wise de Valdez et al., 2011), while Wolbachia-based IIT/SIT combinations suppressed hatching to zero within 12–14 weeks at a 5:1 ratio (Zheng et al., 2019). Notably, even a 1:1 release ratio induced measurable declines, though not full collapse— highlighting a dose-dependent response. Systems like engineered genetic incompatibility (EGI) have reported faster collapse (11–14 weeks at 1:1; Maselko et al., 2020; Upadhyay et al., 2022), which may be attributed to the pre-established transgenic population and the delayed female lethality effect induced by high tetracycline concentrations. In contrast, the antibiotic-free YYAP system achieves suppression without such preconditioning or chemical dependence, underscoring its ecological and operational advantages.

However, the greater innovation of YYAP lies in its replacement mode — a strategy aligned with emerging paradigms of responsible and localized genetic intervention. The >99% lethality of hemizygous offspring when *K*711 males mate with wild-type females establishes a strong underdominance threshold (theoretically ∼2/3 for a single-locus system). Above this threshold, YYAP spreads and stably replaces the population; below it, the element is purged. This threshold property prevents transboundary spread and enables spatial confinement—addressing a central concern in gene drive governance (National Academies of Sciences, Engineering, and Medicine, 2016; James et al., 2023). Our cage trials confirm rapid replacement within four generations under high-release conditions, with populations stabilizing at reduced densities due to the system’s inherent design. Such behavior positions YYAP as an ideal candidate for geographically isolated interventions (e.g., island vector control), where ecological containment is non-negotiable (Esvelt et al., 2014; Champer et al., 2020).

Compared to other threshold-dependent systems—such as UDMEL (Akbari et al., 2013) or EGI (Maselko et al., 2020)—YYAP offers two critical advantages. First, it achieves near-complete postzygotic incompatibility without relying on maternal-effect toxins or sex-specific lethality, enabling deployment across diverse life stages and mating systems. Second, and more fundamentally, YYAP circumvents the dominant resistance pathways that undermine most gene drives. CRISPR-based homing drives are vulnerable to resistance via NHEJ-mediated indels (Unckless et al., 2017), while RNA-targeting systems like *Medea* are thwarted by pre-existing sequence polymorphisms that disrupt small RNA–target pairing. In contrast, YYAP functions through a protein-based toxin – antitoxin mechanism: the toxin cleaves mRNAs nonspecifically, while the antitoxin neutralizes it in YYAP-carrying cells. Resistance would require loss-of-function mutations in the toxin gene—a low-probability event— and even then, alternative toxin – antitoxin pairs can be deployed as backups. Consistent with this, no mutations were detected in toxin or promoter regions among rare surviving hemizygotes, and reconstructed lines maintained full lethality over ten generations.

Admittedly, YYAP’ s high release threshold and 50% embryonic lethality in self-crosses impose production costs that limit its competitiveness with radiation-based SIT in well-established programs. It is not designed for low-cost mass rearing like tetracycline-repressible systems (e.g., fsRIDL). Yet, for species refractory to irradiation—or in contexts where reversibility, ecological containment, and resistance-proof design outweigh deployment speed—YYAP offers a uniquely balanced solution. Critically, both suppression (via male-only releases) and replacement (via balanced releases) can be implemented using the same genetic construct, enabling adaptive management strategies tailored to local ecological and regulatory landscapes.

### Diverse Applications and Broader Implications

The modularity of the YYAP platform, leveraging conserved TA systems, opens a spectrum of applications beyond conventional pest control. It can be adapted for the biocontainment of transgenic organisms, as accidental escapes would be unable to establish populations. In agriculture, it could preserve elite genetic lines by eliminating hybrid offspring from crossbreeding. The observed enlarged larval biomass of hemizygous offspring (Figure 2D) also suggests potential in optimizing resource insects like *Hermetia illucens* for waste processing, maximizing biomass while preventing dispersal (Kaczor et al., 2022).

The YYAP system represents a phenotypic departure from Mendelian inheritance by establishing the first paradigm of obligate co-inheritance of homologous alleles—a synthetic co-dependency that functionally challenges the principle of allelic segregation. While analogous systems exist (e.g., balanced lethals or toxin-antitoxin drives like *Medea*), they permit allele separation in hemizygotes (Wallance, 1994; Akbari et al., 2013), and CRISPR-based drives often operate through homing mechanisms that do not impede allelic disjunction during meiosis (Gantz et al., 2015). YYAP, however, can be viewed as an extreme and generalized form of meiotic drive (Burt and Trivers, 2006). Whereas classic drive elements (e.g., segregation distorters) act by “cheating” during gametogenesis to bias transmission (Lyttle, 1991), YYAP implements a post-meiotic, zygotic enforcement mechanism: although homologs physically separate during meiosis as per Mendel’s First Law, the *phm*-driven toxin-antitoxin configuration ensures survival only of *P/M* transheterozygotes while inducing developmental arrest in *P/+* or *M/+* hemizygotes. This system effectively prohibits the phenotypic manifestation of allelic segregation across generations, enforcing near-100% co-transmission of the allele pair—a drive efficiency surpassing most natural systems.

This synthetic allelic co-dependency establishes an obligate postzygotic barrier with profound evolutionary implications. By inducing >99% larval lethality in hemizygotes from crosses between YYAP carriers and WTs, the system functions as a near-complete barrier to gene flow, creating a reproductively isolated subpopulation. This engineered outcome parallels the effects of supergene complexes in balanced lethal systems and demonstrates how meiotic drive can be transformed into a potent speciation barrier (Phadnis and Orr, 2009; Bladen et al., 2024). However, YYAP achieves near-instantaneous barrier establishment—unlike the gradual accumulation of genetic incompatibilities in chromosomal inversions or the slow evolution of classical meiotic drive systems (Wallace, 1994; Noor et al., 2001; Kirkpatrick and Barton, 2006). Furthermore, its synthetic design circumvents common evolutionary constraints of natural drivers (e.g., fitness costs or suppressor evolution), allowing stable, high-fidelity drive (Greenberg et al., 2025). Consequently, YYAP offers not only a novel pest management tool but also a programmable system to study the mechanisms and dynamics of postzygotic isolation under controlled settings, providing a powerful model for probing how selfish genetic elements catalyze speciation.

### Limitations and Future Directions

Despite its groundbreaking capacity to enforce allelic co-inheritance, the current YYAP system faces practical constraints that merit attention. The larval-stage lethality, while advantageous for density-dependent pest suppression (Juliano, 2007; Deredec et al., 2011; Alphey and Bonsal, 2014), limits efficacy against pests inflicting critical crop damage during early larval stages (e.g., lepidopteran stem borers). This could be mitigated by engineering embryonic lethality using promoters with early developmental activity. Additionally, residual toxicity in carrier males (manifested by a 6-day lifespan reduction) imposes a fitness cost that may compromise population suppression efficacy. This reduced competitiveness necessitates larger release cohorts to achieve target suppression thresholds, thereby escalating operational costs. Future iterations should screen attenuated toxin variants (e.g., MazF mutants with reduced catalytic efficiency) or alternative toxin-antitoxin (TA) pairs sourced from bacterial pangenomic reservoirs (>210,000 loci; Guan et al., 2024), with lower cellular toxicity.

To enhance field applicability, implementing female-specific lethality through sex-spliced effectors (e.g., tra-intron-regulated toxins) combined with fluorescent markers for automated sex-sorting would enable male-only release protocols—eliminating high-damage females while sustaining system propagation via hemizygous males. Beyond TA systems, modular lethality platforms could leverage programmable RNAi (e.g., dsRNA targeting essential genes + rescue cassettes) or CRISPR-ribonucleases for spatiotemporal control. Finally, replacing φC31 integration with CRISPR-Cas9 or piggyBac-Flp/FRT systems would streamline deployment in non-model pests, though rigorous containment testing remains essential prior to ecological use. Collectively, these refinements will transition YYAP from a proof-of-concept toward a versatile toolkit for inheritance-driven biocontrol and speciation research.

It is important to note that while our system achieved effective population modification, the introduction threshold was quite high. Even in ideal form (100% efficiency and no fitness costs), the introduction frequency threshold of this system for population modification is 2/3. Though very confined, this means that it would have trouble spreading from its release region, requiring widespread releases (Champer et al., 2020e). However, by separating each toxin-antidote allele into its own locus, a 2-locus system would be created, reducing the ideal introduction threshold to 27% (though preventing it use in a suppression system) (Magori and Gould 2006; Huang et al., 2009; Huang et al., 2011; Edgington and Alphey 2017; Dhole et al 2018; Champer et al., 2020e; Sánchez et al 2020). This would allow for spread in a continuous region from a single release in just part of the region, greatly increasing drive power while still preserving strong confinement. This property could persist even in the face of moderate drive or cargo fitness costs.

## Acknowledgments

This work was supported by grants from the National Natural Science Foundation of China (No. 31872290). We are grateful for the assistance of all staff members and students in the Laboratory of Pest Control and Biosafety for Staple Crops, Huazhong Agricultural University at Wuhan, Hubei, China.

## Author contributions

Conceptualization, Z. Z. H. and L. Y. F.; methodology, L. Y. F. and Z. Z. H; software, L. Y. F.; formal analysis, L. Y. F and Z. Z. H.; investigation, L. Y. F., Q. Z. Z., X. C. J., H. S. Z., J. J. F., Y. L. P., L. W. P., T. M. M., Y. R. B., and Z. Q. Y.; resources, L. Y. F., Q. Z. Z., X. C. J., H. S. Z., J. J. F., Y. L. P., L. W. P., and Y. R. B.; data curation, L. Y. F., and H. S. Z.; writing – original draft, L. Y. F., and Z. Z. H.; writing – review & editing, Z. Z. H., J. C., and L. Y. F.; visualization, L. Y. F.; supervision, Z. Z. H.,M. W. H., and H. H. X.; funding acquisition, Z. Z. H., M. W. H., and H. H. X.

## Methods

### Plasmid Design and Construction

Representative architectures of the transgenic constructs are shown in Figure S2. The construction process is described here using two type VIII vectors based on the K711 strain as examples; construction of other vector types can be provided upon request. The backbone plasmid pYYAP-basic was derived from pxlBacII and contains an SV40 polyA signal and an attB site. The plasmid was linearized with XhoI and assembled with a *gypsy* insulator fragment to generate pYYAP-basic1.0. A fusion cassette containing *RelB* (*S. pneumoniae*), T2A, *MazFK21A* (*E. coli*), and the reverse-oriented *RelE* (*S. pneumoniae*) was cloned into the KpnI/BamHI sites of pYYAP-basic1.0 to obtain pYYAP-MazFK21A-RelE(Reverse)-2. Next, Opie2, *RelB* (*S. pneumoniae*), T2A, a synonymous *RelB* variant, and the Opie2 polyA signal were PCR-fused and inserted into the XhoI/EcoRI-digested vector to yield pYYAP-MazFK21A-RelE(Reverse)-3. Following Eco91/NotI linearization, the fused cassette containing FRT3, HSTKpolyA, and *mini-white* was cloned in to generate pYYAP-MazFK21A-RelE(Reverse)-4. A separate fusion of OP64polyA, *MazE* (*E. coli*), Acie1, FRT3, phm, and *gypsy* was inserted into the NotI/SpeI sites to produce pYYAP-MazFK21A-RelE(Reverse)-miniwhite.

In parallel, *MazE* (*E. coli*), T2A, *RelE* (*S. pneumoniae*), and reverse-oriented *MazF* (*E. coli*) were PCR-fused and cloned into the Eco91/BamHI-digested pYYAP-basic1.0, generating pYYAP-RelE-MazF(Reverse)-2. Similarly, a fused fragment of Opie2, *MazE* (*E. coli*), T2A, a synonymous *MazE* variant, and Opie2 polyA was inserted into the XhoI/EcoRI-digested vector to obtain pYYAP-RelE-MazF(Reverse)-3. After Eco91/SpeI linearization, a PCR-fused cassette containing FRT5, HSTKpolyA, and *yellow* was inserted, yielding pYYAP-RelE-MazF(Reverse)-4. A final fusion of OP64polyA, *RelB* (*S. pneumoniae*), Acie1, FRT5, phm, and *gypsy* was cloned into the NotI site to produce pYYAP-RelE-MazF(Reverse)-yellow.

Promoter fragments were PCR-amplified from *Drosophila* genomic DNA using the following primer pairs: sry-F/R, phm-F/R, phmΔ228-F/R, mhc-F/R, elav-F/R, dib-F/R, repo-F/R, and haf-F/R. Toxin–antitoxin modules were selected based on previously published studies and the TADB3.0 database (Guan et al., 2024; Li et al., 2025). The sequences of the synonymous *RelB* (*S. pneumoniae*) and *MazE* (*E. coli*) variants are listed in Supplementary Table S6. All constructs were assembled using Gibson enzymatic assembly (Gibson et al., 2009). Descriptions of plasmids and the corresponding transgenic strains are provided in Table S2, and primer information is provided in Table S7.

### Fly Stocks and Rearing Conditions

Flies were maintained at 25 ± 1°C and 60 ± 10% relative humidity. The cornmeal food recipe (per liter) consisted of 6 g agar, 41 g maltose, 40 g sucrose, 9.2 g soy flour, 33 g yellow cornmeal, 25 g yeast, and 10 ml propionic acid. The fly stocks were obtained from the Bloomington Drosophila Stock Center (BDSC) or the Core Facility of *Drosophila* Resource and Technology, CEMCS, CAS. All transgenic flies were generated by φC31-mediated integration, targeting the third chromosome locus *M{3XP3-RFP.attP}ZH-86Fb* (BDSC: 24749). Transgenic flies were balanced over the *y¹w¹*; *Sp/CyO*; *MKRS/TM2* background. Embryo injections were performed by the Core Facility of *Drosophila* Resource and Technology, CEMCS, CAS. Unless otherwise indicated, the *y¹w¹* strain was used as the WT control. Descriptions of fly stocks are provided in Table S1.

### Generation of YYAP Lines

Transgenic lines were initially identified under a fluorescence or stereomicroscope according to their selection markers (*mCherry*/*mini-white*/*yellow*) and subsequently confirmed by PCR genotyping. The crossing strategy used to combine two independent transgenic lines and produce the YYAP line is illustrated in Figure S3. After establishing stable homozygous transgenic lines, these flies were crossed to a line carrying the *FLP* gene (BDSC: 6934). In the F2 generation, individuals carrying both the *FLP* gene and the corresponding selection marker were selected and further crossed. In the F3 generation, individuals carrying the *FLP* gene together with both selection markers were used as parents. After 3–4 days of mating, parental flies were removed, and food vials containing L1–L2 larvae were heat-shocked in a water bath at 37℃ for 30 min per day for 4–5 consecutive days to induce recombination at the FRT sites. Recombination was assessed based on mosaic phenotypes derived from the presence or absence of selection markers. Flies with mosaic phenotypes were further crossed, and offspring exhibiting a white-eye, yellow-body phenotype were preliminarily identified as YYAP lines. PCR genotyping was performed to confirm their genotypes. To ensure genetic homogeneity, offspring from all crosses were isolated at the pupal stage and maintained individually in 2 ml tubes until eclosion. During the establishment of the YYAP line, the number of progeny with different phenotypes was recorded at each stage, and the eclosion rate was calculated as the ratio of pupae that eclosed on a given day to the total pupae collected.

### Molecular Validation of YYAP Cassette Integration and Transcription in *K*711

Genomic DNA was extracted from the transgenic lines P’, M’, or YYAP lines, and PCR was performed with specific primers to amplify the toxin-T2A-antitoxin cassette for genotype validation. The primers used in this study are listed in Table S7. Representative results for lines 2311, 2407, and *K*711 are shown in Figure S3A; other validation data are available upon request. To evaluate toxin expression, total RNA was extracted from the *K*711 line. Reverse transcription was performed using the Takara® RT-PCR kit, with a no-reverse transcriptase control to exclude genomic DNA contamination. Transcriptional validation of *K*711 is shown in Figure S3B. Primer sequences are listed in Table S7. All experiments were performed in three independent biological replicates.

### Viability, Fecundity and Developmental Analyses of YYAP Lines

To test hemizygous lethality of different YYAP lines, three virgin females of the same YYAP line were crossed with three WT males, or three males of the same YYAP line were crossed with three WT virgin females. Homozygous lethality was assessed by crossing males and females of the same YYAP genotype. All crosses were maintained in vials for five days, after which adults were removed to prevent larval overcrowding. The number of adult progeny was scored 15 days after egg laying. To determine the developmental stage at which lethality occurred, we quantified fecundity, hatching rates, and developmental timing in crosses between *K*711 flies and WT flies, with *K*711 serving either as the maternal or paternal genotype. Three replicate crosses were set up, each with 20 virgin females of one genotype (WT or *K*711) and 20 males of the reciprocal genotype, maintained in embryo collection cages with grape juice agar plates. In parallel, three *K*711 intrastrain crosses were performed to assess heterozygote-associated data, and three WT intrastrain crosses were included as controls. The number of eggs laid per plate within a 24 h interval was recorded as the fecundity. For hatching assays, 100 eggs from each plate were monitored for more than 36 h, and the number of unhatched eggs was counted. To evaluate developmental timing and determine the lethal stage, 60–70 eggs from each plate were transferred to food-containing vials and cultured until adulthood. The age and survival status of progeny were examined every 12 h until they either eclosed as adults or died. All experiments were performed in three independent biological replicates.

### Longevity and Survival Analyses of YYAP Lines

To evaluate the survival curves of *K*711 and WT flies, newly eclosed adults were collected and aged in batches of 15–20 flies per vial, separated by sex. Each replicate contained 60–100 flies distributed across four to five vials. Flies were transferred to fresh food every three days, and the number of dead flies was recorded at each transfer. All experiments were performed in three independent biological replicates. Survival data were analyzed using the Lifelines Python package. Survival functions were estimated by the Kaplan–Meier method, differences between groups were assessed by log-rank tests, and 95% confidence intervals were calculated using Greenwood’s formula.

### Male Mating Competitiveness and Sperm displacement Assays

To assess the mating competitiveness of *K*711 males, we compared their ability to inseminate WT females in the presence of WT males. Because *y^1^w^1^* and *K*711 males share the same genetic background, they provide an ideal comparison. In one assay, a single virgin WT female was placed together with one WT male and one *K*711 male for 8 h. After mating, the female was transferred to a new vial for a 4-day oviposition period, and adult progeny were counted 14 days after egg laying. In a second assay, virgin WT females were exposed to varying ratios of WT and *K*711 males under competitive conditions. Each vial contained 20 virgin WT females, and the experiment was carried out in two groups of four vials. In the first group, females were mated with 1–4 WT males, whereas in the second group females were mated with a total of four males, with the WT:*K*711 ratio set at 3:1, 2:2, 1:3, or 0:4. Mating was allowed for 24 h, after which females were transferred to fresh vials for a 24 h oviposition period, and progeny were scored 14 days later. All experiments were performed in three independent biological replicates.

To evaluate the sperm competitiveness of *K*711 males, we first tested their ability to displace sperm previously stored in the seminal receptacle of females. Five-day-old non-virgin WT females, which had already mated with WT males, were individually placed either alone, with three WT males, or with three *K*711 males. Adults were transferred to fresh vials every four days for a total of three sequential egg-laying periods spanning 12 days, and progeny were scored 14 days after each transfer. In a second assay, we assessed the offensive and defensive abilities of *K*711 sperm in sperm displacement. Virgin WT females were paired with either WT or K711 males for an 8 h mating period and then transferred to individual vials for 2 days of oviposition. Females showing evidence of mating were subsequently remated with a male of the alternative genotype for 8 h. After the second mating, females were placed in a new vial for a 2-day oviposition period, followed by transfer to a final vial for an additional 7 days of oviposition. Progeny were scored 14 days after egg laying in both the 2-day and 7-day intervals. For both assays, flies used were 2–5 days old, and successful copulation was confirmed by the presence of fertilized eggs developing into larvae. All experiments were performed in three independent biological replicates.

### Population Suppression and Replacement Cage Trials

In population suppression cage trials, overlapping generations were used to mimic natural population dynamics. At the start of week 1, each cage was seeded with 100 WT flies. To establish suppression conditions, 100 *K*711 males were released weekly into 1:1 suppression cages, and 1000 *K*711 males were released weekly into 10:1 suppression cages, with releases continuing for 8 consecutive weeks. The trials were conducted over 16 weeks in cages of 0.064 m³. Adults were maintained on cotton pads soaked with 15 ml of liquid diet containing 5% yeast extract, 5% sucrose, and 0.6% propionic acid, supplied in 10-cm Petri dishes and replaced twice per week. Once per week, a 250-ml bottle containing 30 ml of cornmeal diet was introduced for 24 h to allow oviposition, after which adults were removed and the bottles sealed. Ten days later, the bottles were opened for 3 days to permit the emerging progeny to mix and mate with the existing population. Population size and genotype composition were monitored weekly using a 24 h trapping assay containing 20 ml of yeast–sucrose solution (0.5% yeast, 2.5% sucrose). To measure the reproductive output of WT females, random females from traps were isolated once per week into three vials for a 3-day oviposition period before being returned to cages. Progeny emerging 14 days later were quantified. All groups were set up in triplicate.

Population replacement cage trials were conducted with non-overlapping generations using 250-ml cornmeal bottles. Each cage was initially seeded with 15 WT females and 15 WT males. In replacement cages, additional *K*711 flies were introduced at ratios of 1:1 (15 *K*711 males and 15 *K*711 females), 5:1 (75 *K*711 males and 75 *K*711 females), or 10:1 (150 *K*711 males and 150 *K*711 females). Control cages without K711 flies were included, and all groups were set up in triplicate. Each generation was allowed to mate and oviposit for 72 h, after which adults were removed. Eleven days later, progeny were screened for genotype by fluorescence and used to establish the next generation. This process was repeated for 10 generations.

### Mathematical modeling

We simulated YYAP-based suppression and replacement strategies in *A. aegypti* populations using the MGDrivE framework (Sánchez et al., 2019). MGDrivE is designed to study the dynamics of gene drive systems and their spatial spread in mosquito populations. The model incorporates daily time steps to capture life stages (egg, larva, pupa, and adult) and includes overlapping generations with explicit mating structure. Males are assumed to mate multiple times throughout life, while females mate once upon emergence and retain the genetic material of their mate for their entire adult lifespan. Larval mortality is density-dependent following the functional form described in Deredec et al. (2011). The inheritance of YYAP was modeled within the Mendelian inheritance module of MGDrivE. The initial population was set at 10,000 adult females. Two simulation scenarios were designed to reflect distinct control strategies. For population suppression, YYAP males were released weekly over a 6-month period to evaluate their impact on reducing population size. For population replacement, one or two combined releases of both males and females were performed to assess their potential to drive substitution. All simulations were conducted within the stochastic version of the MGDrivE framework and repeated 1,000 times. Life-history parameters and intervention settings are provided in Table S5.

### Data analysis, statistics, and visualization

Data were analyzed using IBM SPSS Statistics v21. Statistical significance thresholds were set as follows: *p* < 0.05 (*), *p* < 0.01 (**), and *p* < 0.001 (***). Figures were generated with GraphPad Prism 8 and edited in Adobe Illustrator to meet publisher requirements. For lethality assays (Figures 2C, 3D, 3E and S6), comparisons were performed using *t* tests assuming unequal variance. Figure S8A was analyzed by Student’s *t* test, whereas Figure S8B used Kruskal–Wallis one-way ANOVA with multiple comparisons. For Figure 4C, a Mann–Whitney test was applied. Emergence rates in Figure S6 were analyzed by Welch’s ANOVA with Tamhane’s multiple comparisons. Sex ratio data were analyzed by Pearson’s chi-squared test. Figures 3A and S7B used one-way ANOVA followed by Tukey’s post hoc test, while Figure S7C employed Welch’s ANOVA with Tamhane’s multiple comparisons. Survival curve data in Figures 3B and 3C were analyzed using the Lifelines Python package. Survival functions were estimated by Kaplan–Meier, with differences assessed by log-rank test and 95% confidence intervals calculated using Greenwood’s method. No blinding was performed during experimentation or outcome assessment.

## Supplementary Information

**Supplementary Figure 1.**
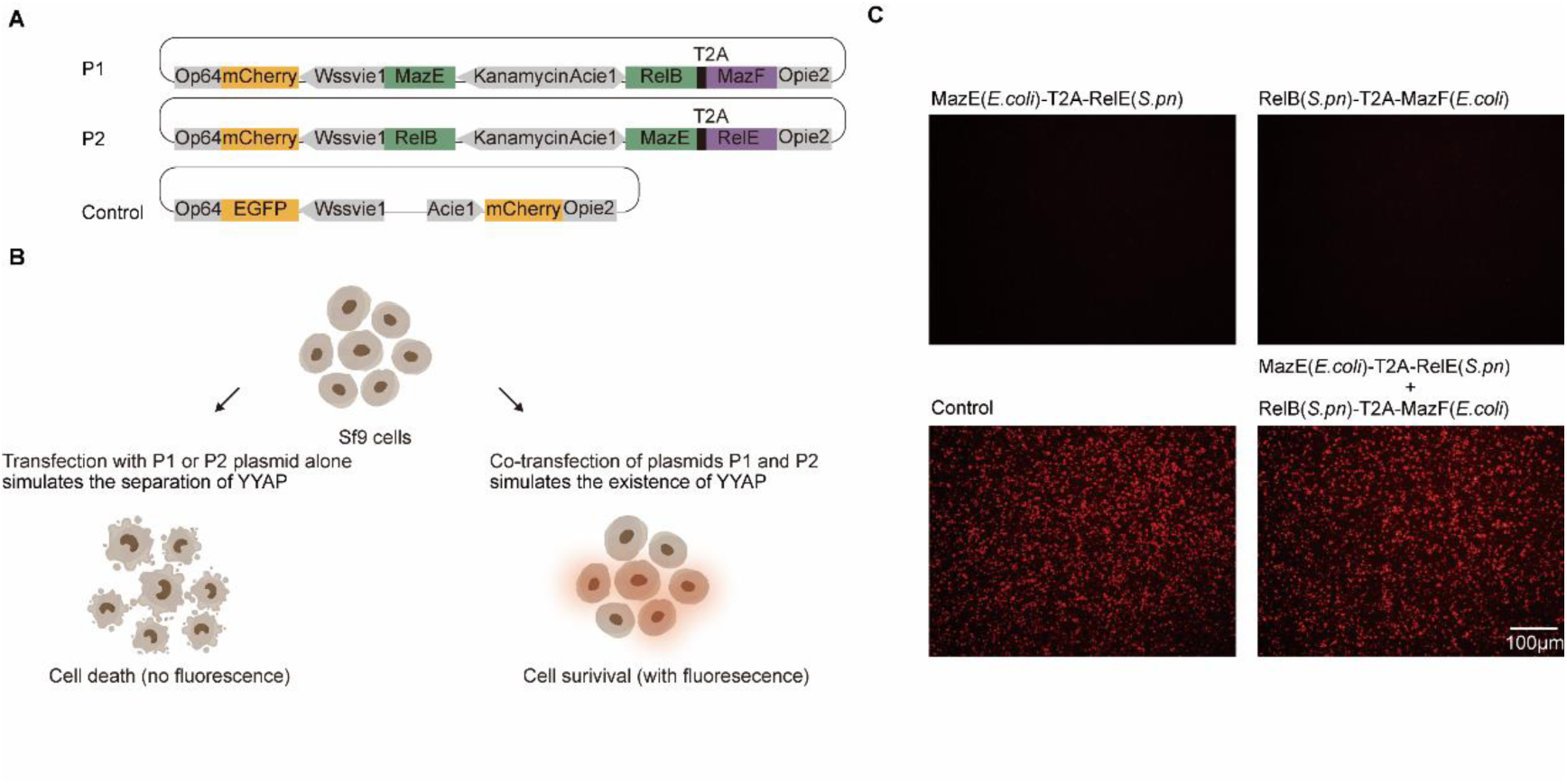
Schematic illustration of YYAP cell experiments. (A) Plasmid constructs. Plasmid P1 simulates allele P and encodes the toxin MazF (*E. coli*) together with the antitoxin RelB (*S. pneumoniae*). Plasmid P2 simulates allele M and encodes the toxin RelE (*S. pneumoniae*) together with the antitoxin MazE (*E. coli*). Toxin–antitoxin pairs (antitoxin B-toxin A or antitoxin B-toxin A) are linked via a T2A peptide and co-expressed under the control of the Acie1 promoter. An Wssvie1-driven *mCherry* reporter serves as a visual indicator of toxicity. (B) Cell-based assay. Co-transfection of both plasmids into Sf9 cells simulates the presence of YYAP, allowing mutual neutralization between toxins and antitoxins. Under this condition, cells survive normally, and the number of red fluorescent cells is not significantly different from that of controls. In contrast, transfection with a single plasmid expressing either antitoxin B–toxin A or antitoxin A–toxin B simulates YYAP segregation, where the absence of the corresponding antitoxin leads to toxin-induced cell death and a reduction in red fluorescent cells. (C) Validation of the YYAP concept in Sf9 cells. Cells were transfected with pmCherry-MazE (*E. coli*)-T2A-RelE (*S. pneumoniae*), pmCherry-RelB (*S. pneumoniae*)-T2A-MazF (*E. coli*), or the control vector pmCherry-EGFP, in the indicated combinations. Images were acquired 3 days post-transfection. Scale bars, 100 μm Abbreviations: Opie2, *Orgyia pseudotsugata multicapsid nucleopolyhedrovirus* (OpMNPV) immediate-early 2 (IE2) polyA signal; Wssvie1, *White spot syndrome virus* (WSSV) immediate-early 1 (IE1) promoter; Acie1, *Autographa californica multiple nucleopolyhedrovirus* (AcMNPV) immediate-early 1 (IE1) promoter; Op64, OpMNPV major envelope glycoprotein (GP64) polyA signal; Kanamycin, kanamycin resistance promoter.

**Supplementary Figure 2.**
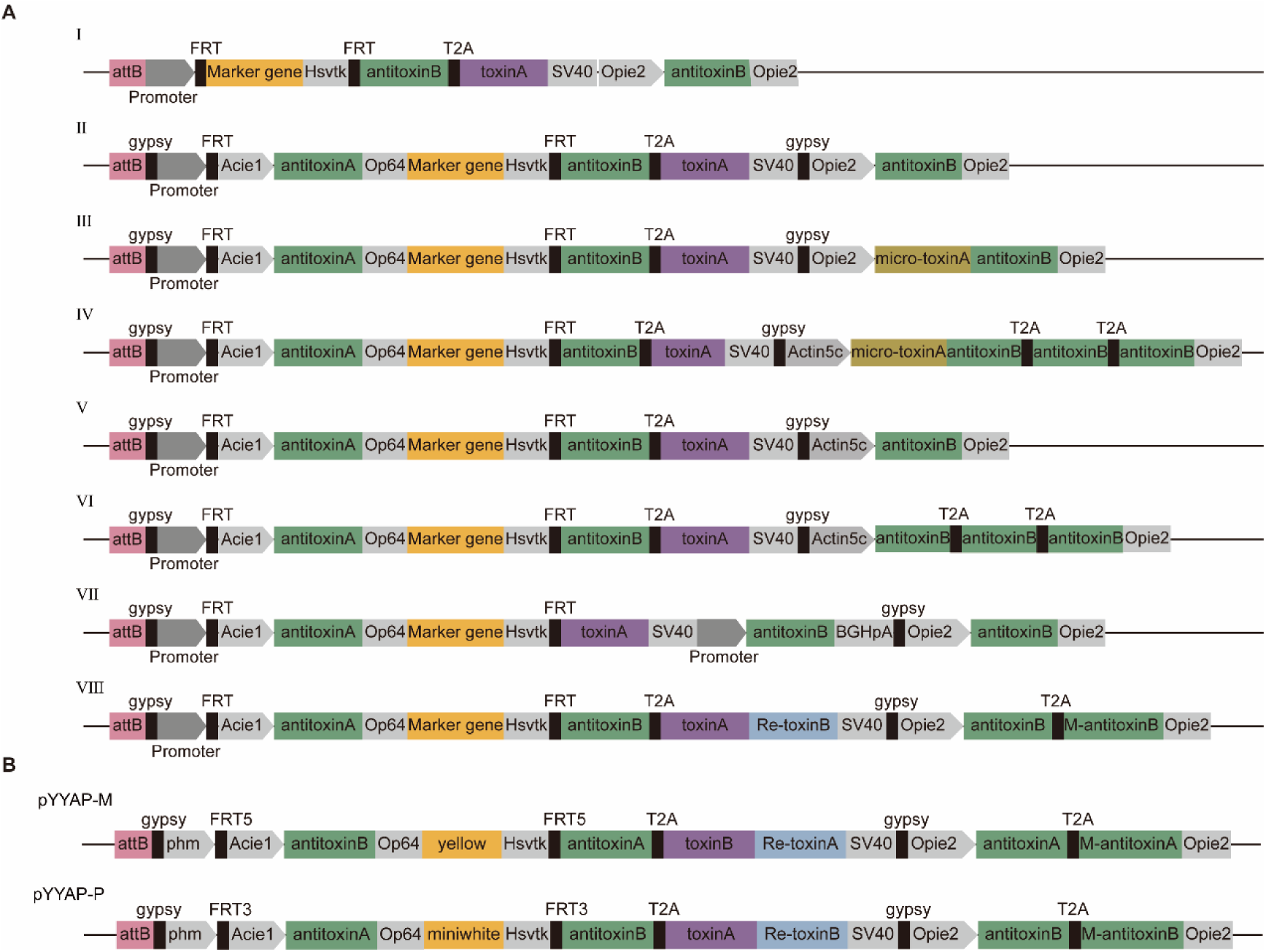
Schematic diagrams of all YYAP plasmid constructs. (A) All constructs contain an attB site for site-specific recombination mediated by φC31 recombinase. Type I: The initial design in which the promoter::antitoxin-T2A-toxin cassette is interrupted by an FRT-Marker-FRT sequence. The Opie2::antitoxin-Opie2p cassette provides constitutive antitoxin expression. Type II: Built on Type I by adding gypsy insulator elements and the Acie1::antitoxin-Op64 cassette, which provides constitutive antitoxin expression to neutralize low-level toxin activity arising from transcriptional readthrough, thereby improving containment. Type III: Incorporates microRNA fragments into the constitutively expressed antitoxin cassette to specifically target RNA of highly virulent toxins. Type IV: Adds two tandem copies of the antitoxin fragment into the constitutive antitoxin cassette and replaces the Opie2 promoter with the endogenous, ubiquitously active Actin5c promoter of *D. melanogaster*. Type V: Derived from Type II, but replaces the constitutive antitoxin promoter with the Actin5c promoter. Type VI: A variant of Type IV lacking the microRNA cassette. Type VII: Splits the original promoter::antitoxin-T2A-toxin cassette into two independent expression cassettes: promoter::toxin-SV40 and promoter::antitoxin-BGHpA. Type VIII: An upgraded version of Type II that introduces an antisense RNA cassette (Re-toxin) downstream of the toxin, targeting the toxin on another chromosome. Additionally, a synonymous mutant antitoxin is incorporated into the constitutive antitoxin cassette. (B) Example of a specific component assembly based on the type VIII genetic construct described in this study. Abbreviations: Hsvtk, *Herpes simplex virus thymidine kinase* (HSV-TK) polyA signal; SV40, *Simian vacuolating virus 40* (SV40) polyA signal; BGHpA, *Bovine Growth Hormone* (BGH) polyA signal.

**Supplementary Figure 3.**
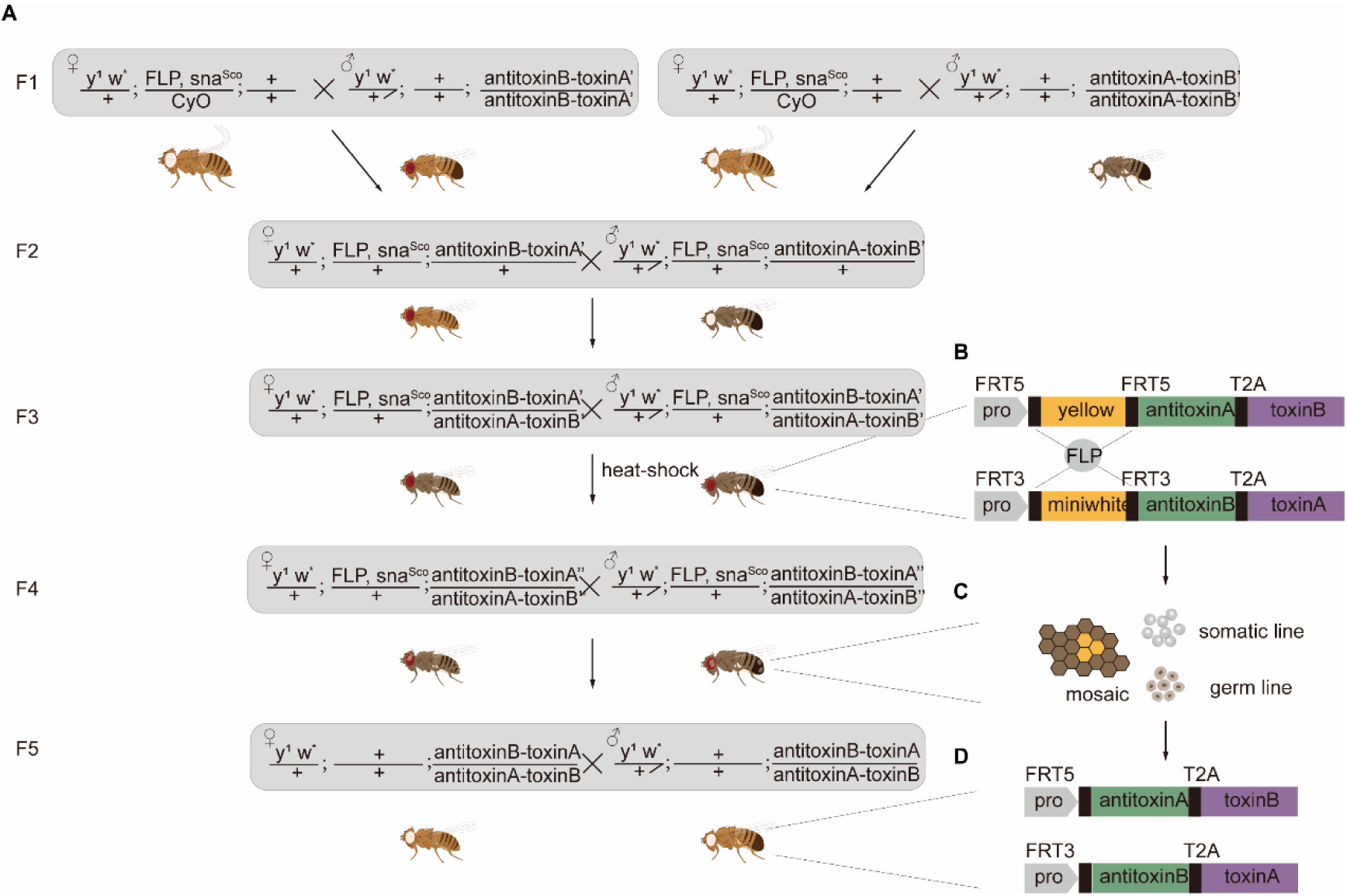
Crossing strategy for generating YYAP fly strains. (A) Genotypes and sexes of flies used in the crosses required to establish YYAP strains are shown. Each cross is indexed with the prefix “F”. “×” indicates mating between parents, while “+” denotes the WT allele. Heat shock treatment was performed by placing vials containing L1 and L2 larvae in a 37°C water bath for 30 min per day over five consecutive days, thereby inducing expression of FLP recombinase and enabling recombination between FRT sites. “Antitoxin–toxin′” indicates the state without induced recombination. “Antitoxin–toxin′′” denotes a mosaic state in which some cells have undergone recombination while others have not. “Antitoxin–toxin” refers to the YYAP strain derived from recombinant germ cells, at which point all cells uniformly express the antitoxin–toxin construct. (B-D) Illustration of the recombination process. (B) FLP recombinase recognizes FRT sites and initiates recombination. (C) Partial recombination in somatic and germ cells generates mosaic flies. (D) Stable YYAP strains are established following germline recombination.

**Supplementary Figure 4.**
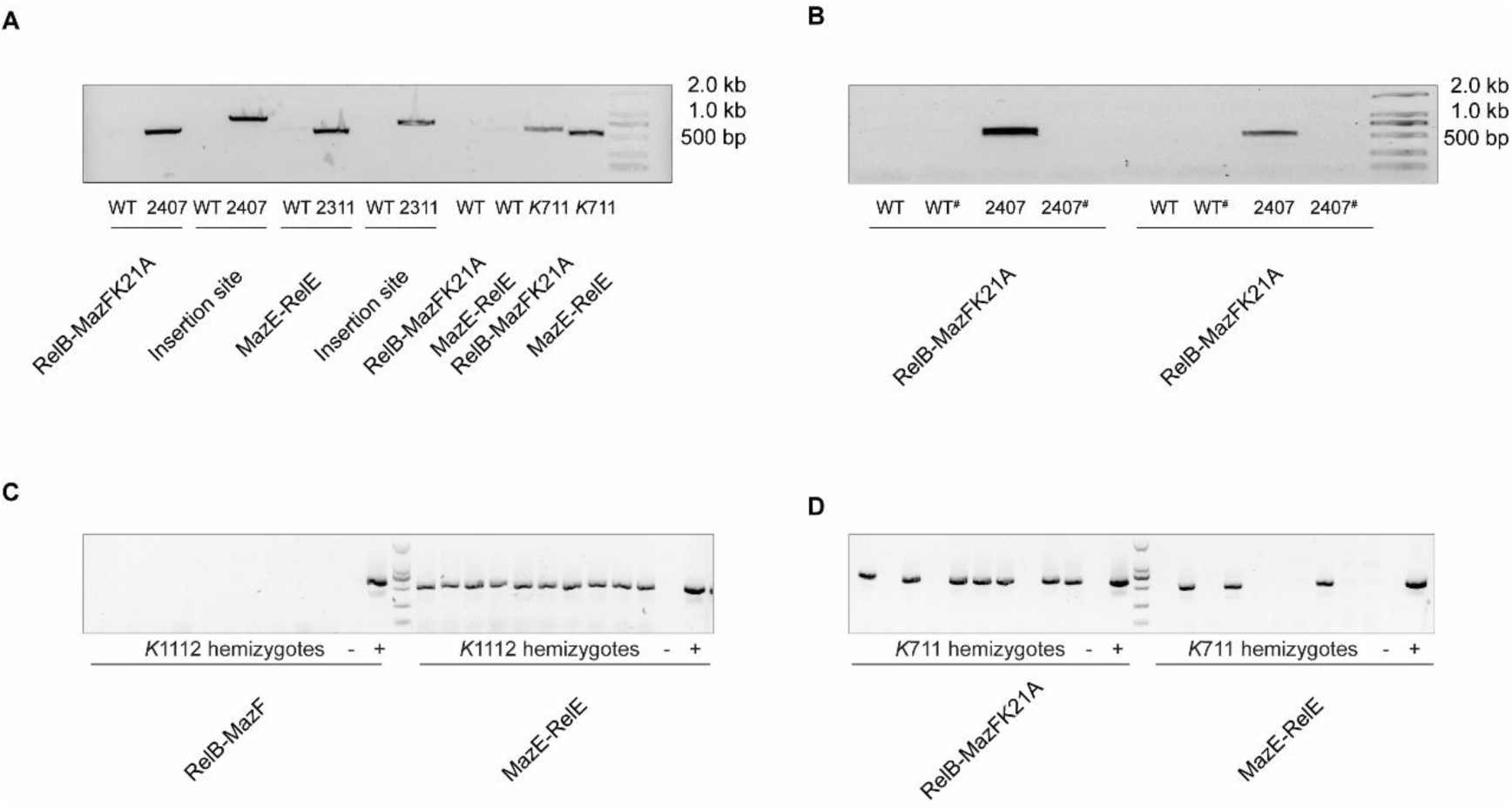
Genotype and transcription verification of representative transgenic strains. (A) PCR was performed using one primer upstream of the insertion site and one primer within the inserted fragment to confirm the presence of the antitoxin-toxin construct and to verify its correct integration at the 86F locus. Genotype verification of other transgenic strains or YYAP strains can be provided upon request. (B) RT-PCR was used to assess transcription of the antitoxin–toxin construct. WT refers to the background strain BDSC:24749 used for microinjection. # indicates the no reverse transcriptase (–RT) control. The sequences of the detection primers are listed in Table S6. (C–D) Genotype identification of surviving hemizygotes. (C) *K*1112 hemizygotes; (D) *K*711 hemizygotes. – indicates the negative control; + indicates the positive control.

**Supplementary Figure 5.**
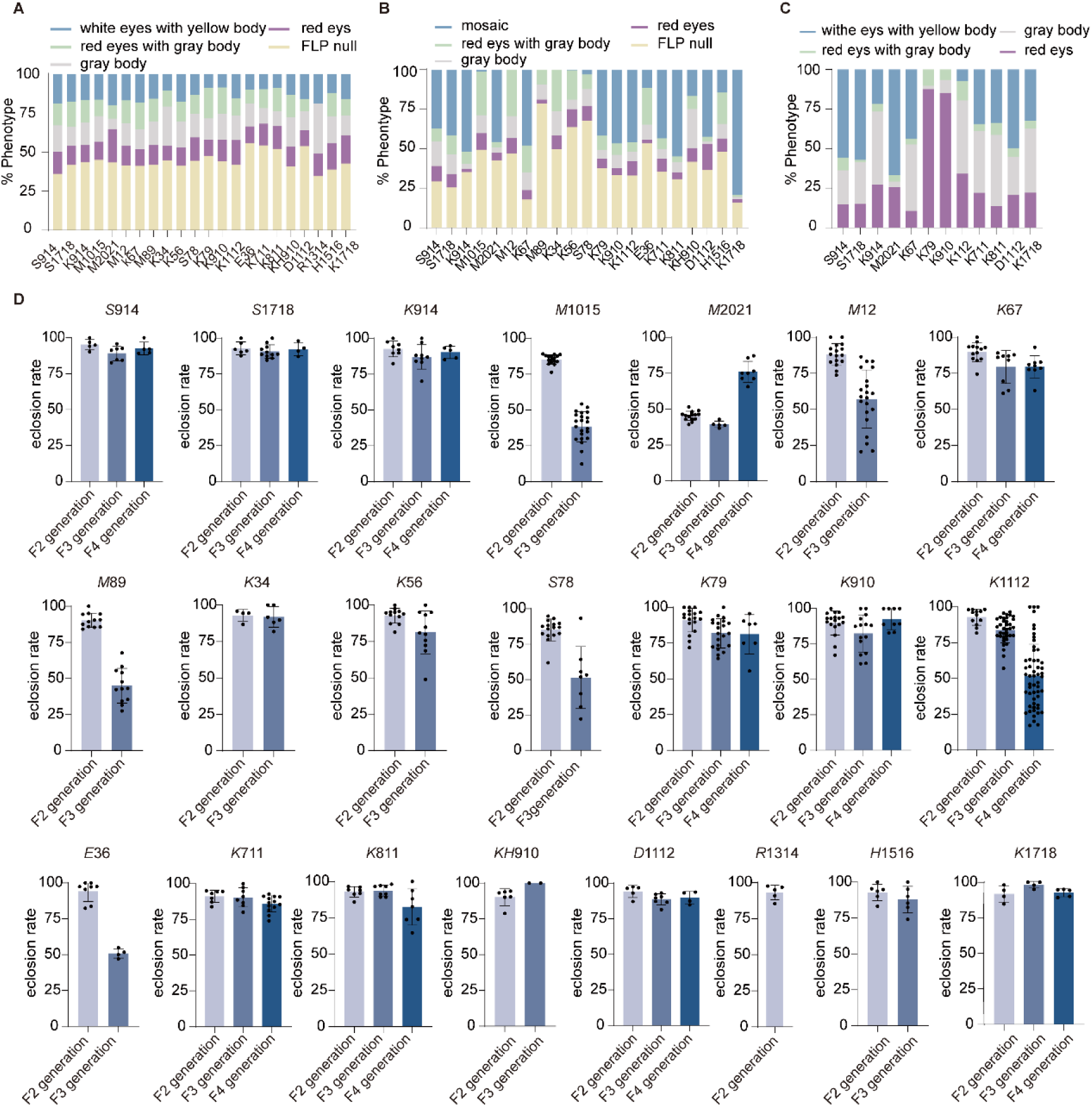
Phenotypic distribution and eclosion rates during the construction of YYAP fly strains. (A) Offspring phenotypic distribution after mating of F2 generation parents, shown as stacked bar charts. FLP null: offspring lacking the *FLP* gene (yellow). White eyes with yellow body: regarded as WT (blue). Red eyes: offspring carrying one toxin–antitoxin construct (dark purple). Gray body: offspring carrying another toxin–antitoxin construct (light purple). Red eyes with gray body: offspring carrying both toxin–antitoxin constructs (green). Representative counts: *S*914 *n* = 164; *S*1718 *n* = 187; *K*914 *n* = 260; *M*1015 *n* = 583; *M*2021 *n* = 346; *M*12 *n* = 469; *K*67 *n* = 552; *M*89 *n* = 412; *K*34 *n* = 235; *K*56 *n* = 578; *K*78 *n* = 669; *K*79 *n* = 703; *K*910 *n* = 924; *K*1112 *n* = 603; *E*36 *n* = 229; *K*711 *n* = 381; *K*811 *n* = 435; *KH*910 *n* = 279; *D*1112 *n* = 275; *R*1314 *n* = 176; *H*1516 *n* = 281; *K*1718 *n* = 199. (B) Offspring phenotypic distribution after mating of F3 generation parents. FLP null: offspring lacking the *FLP* gene (yellow). Mosaic: offspring with somatic recombination and possible germline recombination (blue). Red eyes: offspring carrying one toxin–antitoxin construct (dark purple). Gray body: offspring carrying another toxin–antitoxin construct (light purple). Red eyes with gray body: offspring carrying both toxin–antitoxin constructs (green). Representative counts: *S*914 *n* = 158; *S*1718 *n* = 445; *K*914 *n* = 307; *M*1015 *n* = 592; *M*2021 *n* = 59; *M*12 *n* = 500; *K*67 *n* = 337; *M*89 *n* = 247; *K*34 *n* = 125; *K*56 *n* = 619; *K*78 *n* = 65; *K*79 *n* = 470; *K*910 *n* = 478; *K1112 n* = 1707; *E*36 *n* = 43; *K*711 *n* = 402; *K*811 *n* = 593; *KH*910 *n* = 24; *D*1112 *n* = 355; *H*1516 *n* = 96; *K*1718 *n* = 134. (C) Offspring phenotypic distribution after mating of F4 generation parents. White eyes with yellow body: offspring derived from germline recombination, in which both toxin–antitoxin constructs are expressed (blue). Red eyes: offspring carrying one toxin–antitoxin construct (dark purple). Gray body: offspring carrying another toxin–antitoxin construct (light purple). Red eyes with gray body: offspring carrying both toxin–antitoxin constructs (green). Representative counts: *S*914 *n* = 185; *S*1718 *n* = 133; *K*914 *n* = 345; *M*2021 *n* = 230; *K*67 *n* = 1198; *K*79 *n* = 185; *K*910 *n* = 252; *K*1112 *n* = 3436; *K*711 *n* = 1019; *K*811 *n* = 633; *D*1112 *n* = 134; *K*1718 *n* = 176. (D) Eclosion rates of offspring from the F2, F3, and F4 generations during the construction of different YYAP strains. Representative sample sizes: *S*914 F2 *N* = 5, *n* = 183; F3 *N* = 7, *n* = 181; F4 *N* = 5, *n* = 239. *S*1718 F2 *N* = 6, *n* = 202; F3 *N* = 11, *n* = 459; F4 *N* = 4, *n* = 166. *K*914 F2 *N* = 8, *n* = 326; F3 *N* = 9, *n* = 363; F4 *N* = 5, *n* = 319. *M*1015 F2 *N* = 16, *n* = 592; F3 *N* = 23, *n* = 1630. *M*2021 F2 *N* = 14, *n* = 823; F3 *N* = 5, *n* = 164; F4 *N* = 8, *n* = 324. *M*12 F2 *N* = 16, *n* = 648; F3 *N* = 21, *n* = 986. *K*67 F2 *N* = 13, *n* = 621; F3 *N* = 9, *n* = 379; F4 *N* = 9, *n* = 1596. *M*89 F2 *N* = 12, *n* = 553; F3 *N* = 12, *n* = 438. *K*34 F2 *N* = 4, *n* = 290; F3 *N* = 6, *n* = 146. *K*56 F2 N=12, *n* = 641; F3 *N* = 11, *n* = 559. *K*78 F2 *N* = 16, *n* = 805; F3 *N* = 8, *n* = 159. *K*79 F2 *N* = 18, *n* = 570; F3 *N* = 21, *n* = 684; F4 *N* = 7, *n* = 137. *K*910 F2 *N* = 18, *n* = 757; F3 *N* = 15, *n* = 505; F4 *N* = 9, *n* = 237. *K*1112 F2 *N* = 12, *n* = 620; F3 *N* = 36, *n* = 2353; F4 *N* = 89, *n* = 7289. *E*36 F2 *N* = 8, *n* = 319; F3 *N* = 4, *n* = 89. *K*711 F2 *N* = 7, *n* = 497; F3 *N* = 8, *n* = 394; F4 *N* = 13, *n* = 1182. *K*811 F2 *N* = 7, *n* = 534; F3 *N* = 8, *n* = 694; F4 *N* = 7, *n* = 850. *KH*910 F2 *N* = 6, *n* = 369; F3 *N* = 2, *n* = 28. *D*1112 F2 N=5, *n* = 226; F3 *N* = 7, *n* = 378; F4 *N* = 4, *n* = 186. *R*1314 F2 *N* = 5, *n* = 206. *H*1516 F2 *N* = 6, *n* = 344; F3 *N* = 6, *n* = 136. *K*1718 F2 *N* = 4, *n* = 216; F3 *N* = 4, *n* = 131; F4 *N* = 5, *n* = 215.

**Supplementary Figure 6.**
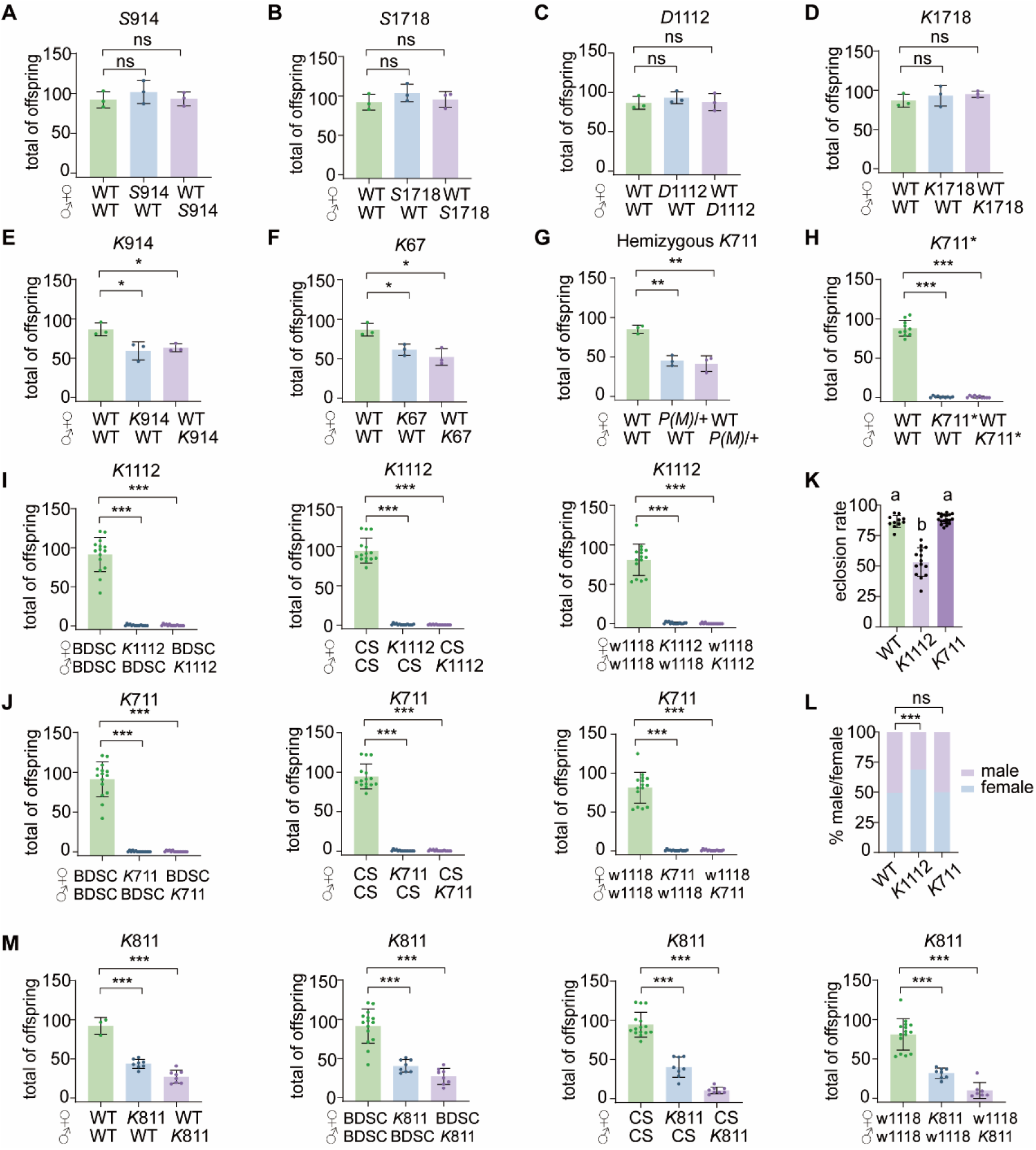
Hemizygous lethality of YYAP. (A-F) Offspring number data for *S*914, *S*1718, *D*1112, *K*1718, *K*914, and *K*67 strains (n = 3). Each panel shows the number of surviving adult offspring from the indicated cross combination. (G) Surviving adult offspring number from crosses between hemizygous *K*711 (genotype: *P/+* or *M/+*) and WT (*n* = 3). (H) Surviving adult offspring number from crosses between *K*711* (genotype: *P/M*) and WT (*n* = 10). Here, *K*711* denotes the *K*711 (genotype: *P/M*) strain re-derived by crossing hemizygous K711 (genotype: *P/+*) with hemizygous *K*711 (genotype: *M/+*). (I) Surviving adult offspring number from crosses between *K*1112 and BDSC:24749/CS/w1118 (*n* = 15). (J) Surviving adult offspring number from crosses between *K*711 and BDSC:24749/CS/w1118 (*n* = 15). (K) Eclosion rates for *K*1112 and *K*711 strains. Welch’s ANOVA followed by Tamhane’s multiple comparisons test was performed across the three strains: WT (*N* = 11, *n* = 1177), *K*1112 (*N* = 14, *n* = 1177), and *K*711 (*N* = 19, *n* = 1497). (L) Male-to-female ratios for *K*1112 and *K*711 strains. Each YYAP strain was compared to the control group using Pearson’s chi-square test: WT (*n* = 717), *K*1112 (*n* = 511), *K*711 (*n* = 1007). (M) Surviving adult offspring number from crosses between *K*811 and BDSC:24749/CS/w1118. For the WT self-cross group, *n* = 3; for all other cross groups, *n* = 8.Statistical significance: ns, not significant; * *p* < 0.05, ** *p* < 0.01, *** *p* < 0.001. Letters in panel (K) indicate significance at α = 0.05.

**Supplementary Figure 7.**
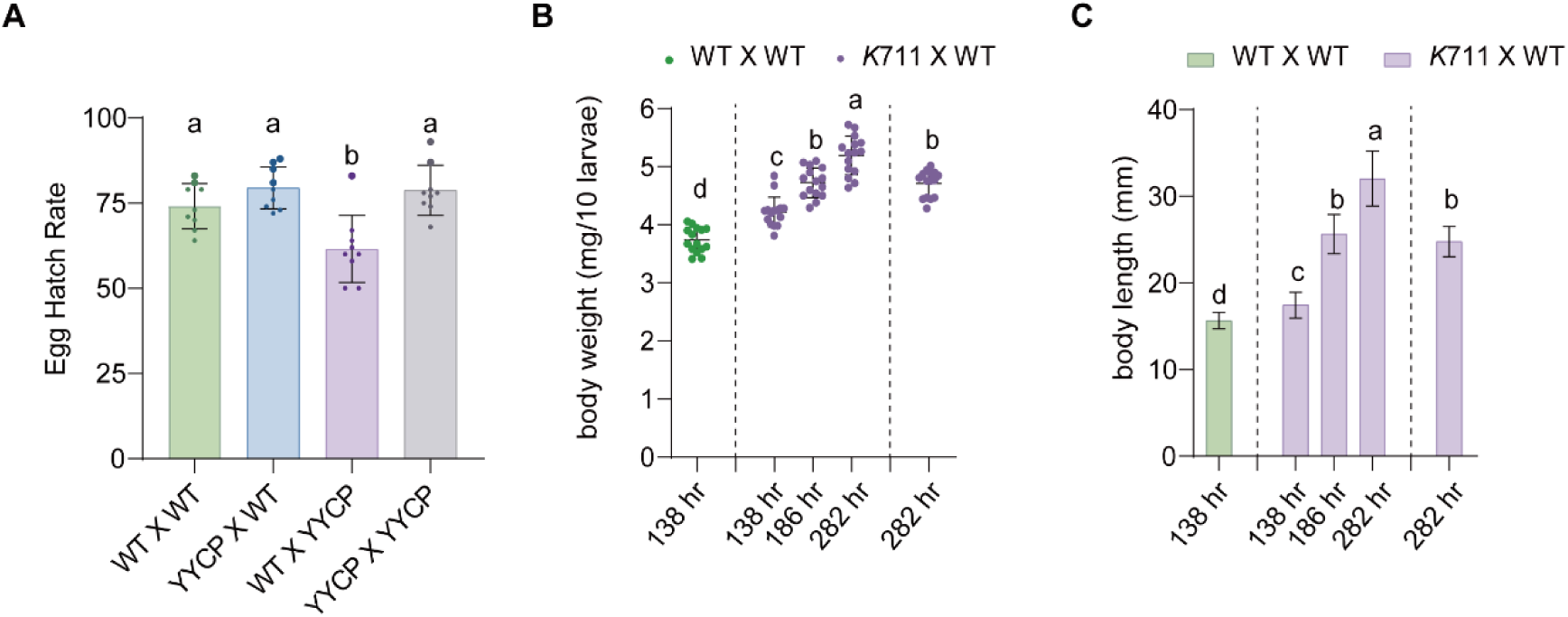
Developmental, and lifespan phenotypes of *K*711. (A) Embryonic hatching rates (%) across four mating groups. A one-way ANOVA followed by Tukey’s multiple comparisons was performed on the hatching rates (*n* = 9). (B and C) Body weight (B) and body length (C) of hemizygous offspring *(P/+* or *M/+*) from *K*711 × WT crosses at three time points (*n* = 15). At 282 hr, two larval phenotypes were observed: larvae continuing to feed within the food (left of the dashed line) and larvae that had climbed onto the vial wall and ceased feeding (right of the dashed line). Body weight data were analyzed using one-way ANOVA with Tukey’s multiple comparisons, while body length was analyzed using Welch’s ANOVA with Tamhane’s multiple comparisons. Letters in panels indicate significance at α = 0.05.

**Supplementary Figure 8.**
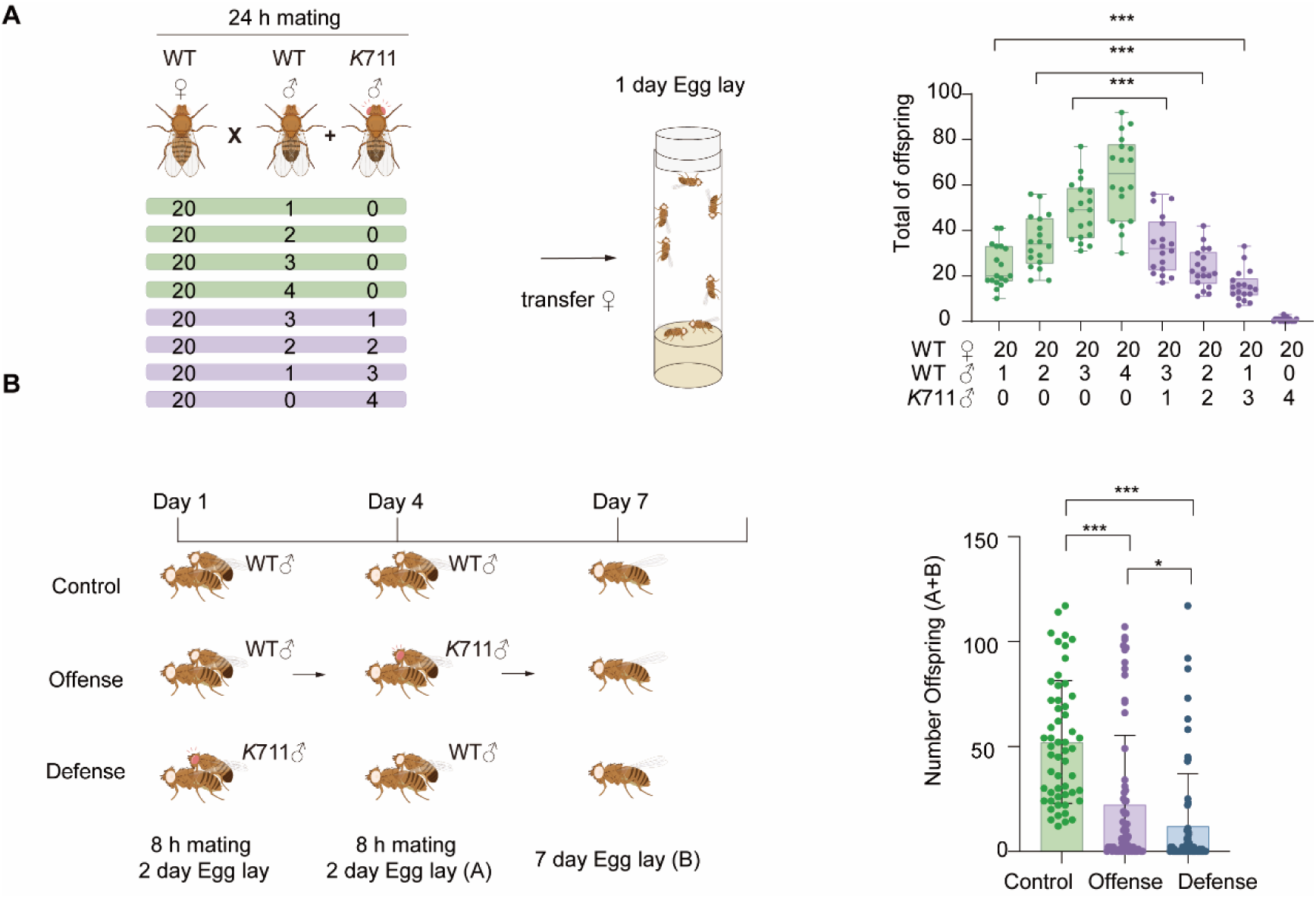
Competitiveness and sperm displacement of *K*711 males. (A) Schematic diagram (left panel) and results (right panel) of the mixed-male mating assay with excess females. Box plots show the median (center line) and interquartile range (box boundaries). Student’s t-tests were performed between experiments with identical numbers of WT males (*n* = 18 per group). (B) Experimental design (left panel) and results (right panel) of offensive and defensive sperm displacement assays. Bar graphs show the number of offspring produced by: one WT female mated to a WT male followed by remating with a second WT male (green, *n* = 55); one WT female mated to a WT male followed by remating with a *K*711 male (purple, *n* = 69); or one WT female mated to a *K*711 male followed by remating with a WT male (blue, *n* = 65). Statistical comparisons among the three groups were performed using Kruskal–Wallis one-way ANOVA with multiple comparisons. Statistical significance: * *p* < 0.05, ** *p* < 0.01, *** *p* < 0.001.

**Supplementary Figure 9.**
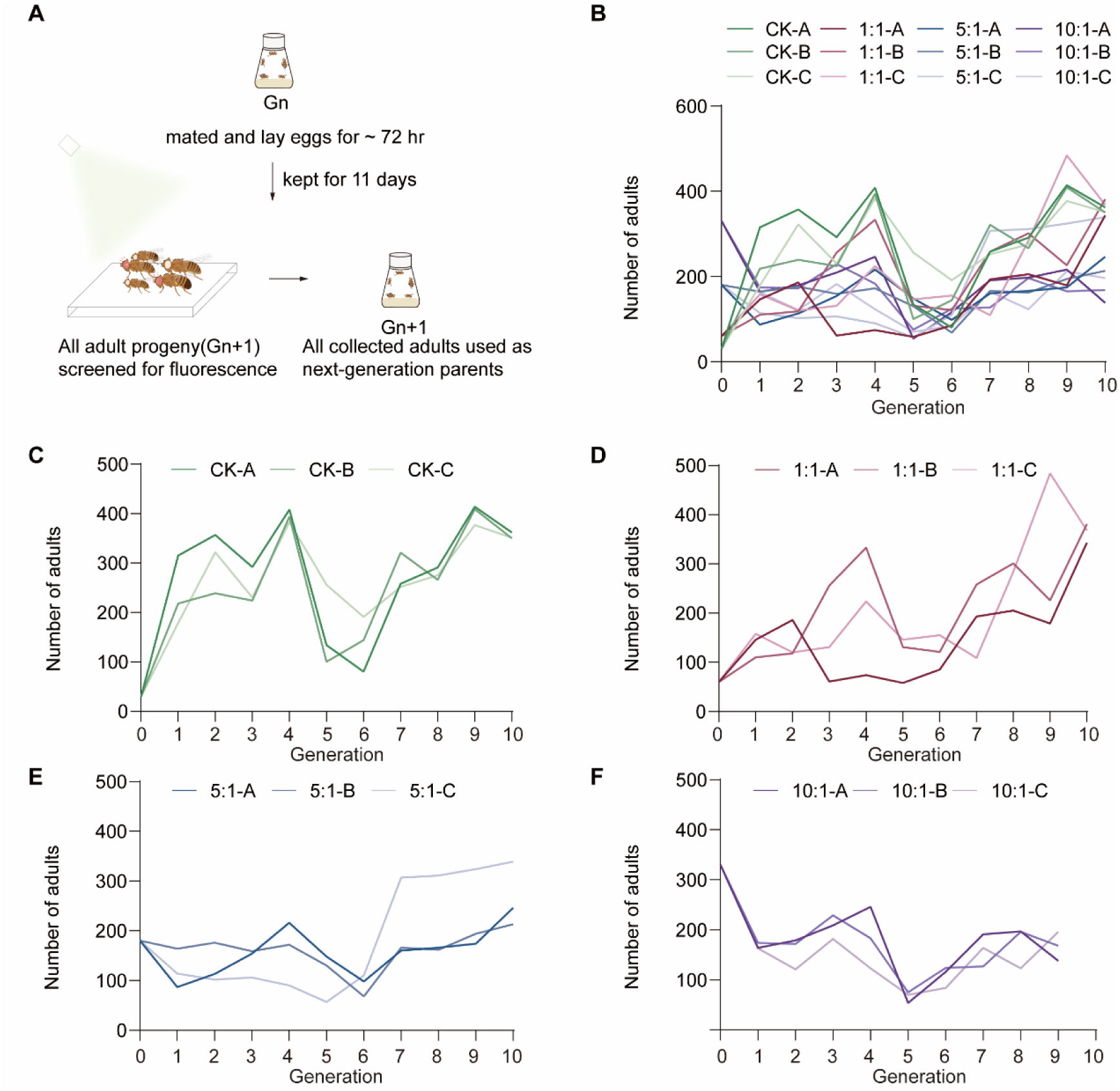
Cage design and population dynamics in replacement experiments. (A) Experimental workflow for population replacement. Parental flies (Gₙ) were allowed to mate and oviposit for ∼72 hr before removal. After 11 days, all emerging adult progeny (Gₙ₊₁) were collected and screened for red fluorescent protein (RFP) expression in the eyes under a fluorescence microscope to assess population composition. All collected adults were then used to seed the next generation, and the process was repeated for 10 generations. (B–F) Population size dynamics in non-overlapping generation small-cage replacement experiments with different release ratios of *K*711. Each cage was seeded with 15 WT females and 15 WT males, along with additional *K*711 females and males to achieve release ratios of 1:1, 5:1, or 10:1 (15♀ + 15♂, 75♀ + 75♂, or 150♀ + 150♂ *K*711 flies, respectively). Offspring in subsequent generations were scored as RFP-positive (due to a 3×P3::RFP cassette upstream of the insertion site) or WT. The total number of adults in each cage (y-axis) was recorded for every generation (x-axis). (B) Combined plot of offspring number dynamics from individual cages. (C) Offspring number dynamics in WT cages. (D) Offspring number dynamics in 1:1 treatment cages. (E) Offspring number dynamics in 5:1 treatment cages. (F) Offspring number dynamics in 10:1 treatment cages.

**Supplementary Figure 10.**
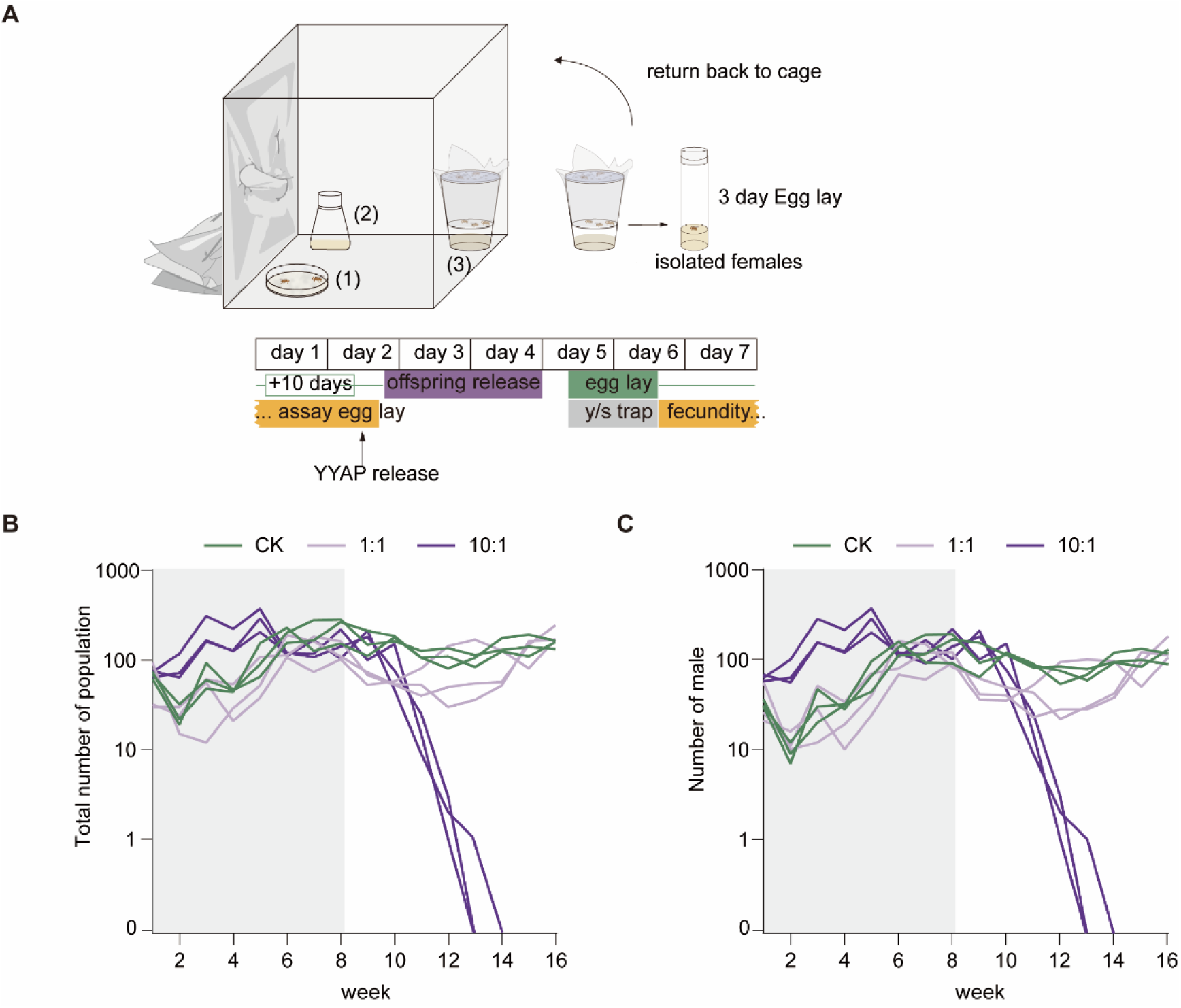
Schematic diagram of the population suppression assay. (A) Cage contains adult feeding apparatus (1), reproduction bottle (2), trapping devices (3) and fecundity assay vials. The weekly experimental timeline is shown below. The trials were conducted over 16 weeks in cages of 0.064 m³. Adults were maintained on cotton pads soaked with 15 ml of liquid diet containing 5% yeast extract, 5% sucrose, and 0.6% propionic acid, supplied in 10-cm Petri dishes (1) and replaced twice per week. Once per week, a 250-ml bottle containing 30 ml of cornmeal diet (2) was introduced for 24 h to allow oviposition, after which adults were removed and the bottles sealed. Ten days later, the bottles were opened for 3 days to permit the emerging progeny to mix and mate with the existing population. Population size and genotype composition were monitored weekly using a 24 h trapping assay containing 20 ml of yeast–sucrose solution (0.5% yeast, 2.5% sucrose) (3). To measure the reproductive output of WT females, random females from traps were isolated once per week into three vials for a 3-day oviposition period before being returned to cages. (B-C) Population dynamics measured from trapping devices. (B) Total population counts. (C) Male counts. Light gray shading indicates weekly additional *K*711 releases through week 8. In control cages (green lines), the population increased gradually and plateaued by week 8. In 1:1 suppression cages (light purple lines), populations increased slowly until week 6, then began to decline; in one cage, recovery began at week 10, while in the other two cages, populations started to rise again by week 12. In 10:1 suppression cages (dark purple lines), the population initially exceeded control levels due to the high release ratio of *K*711 males, followed by population collapse within 5-6 weeks after the releases ceased.

**Supplementary Figure 11.**
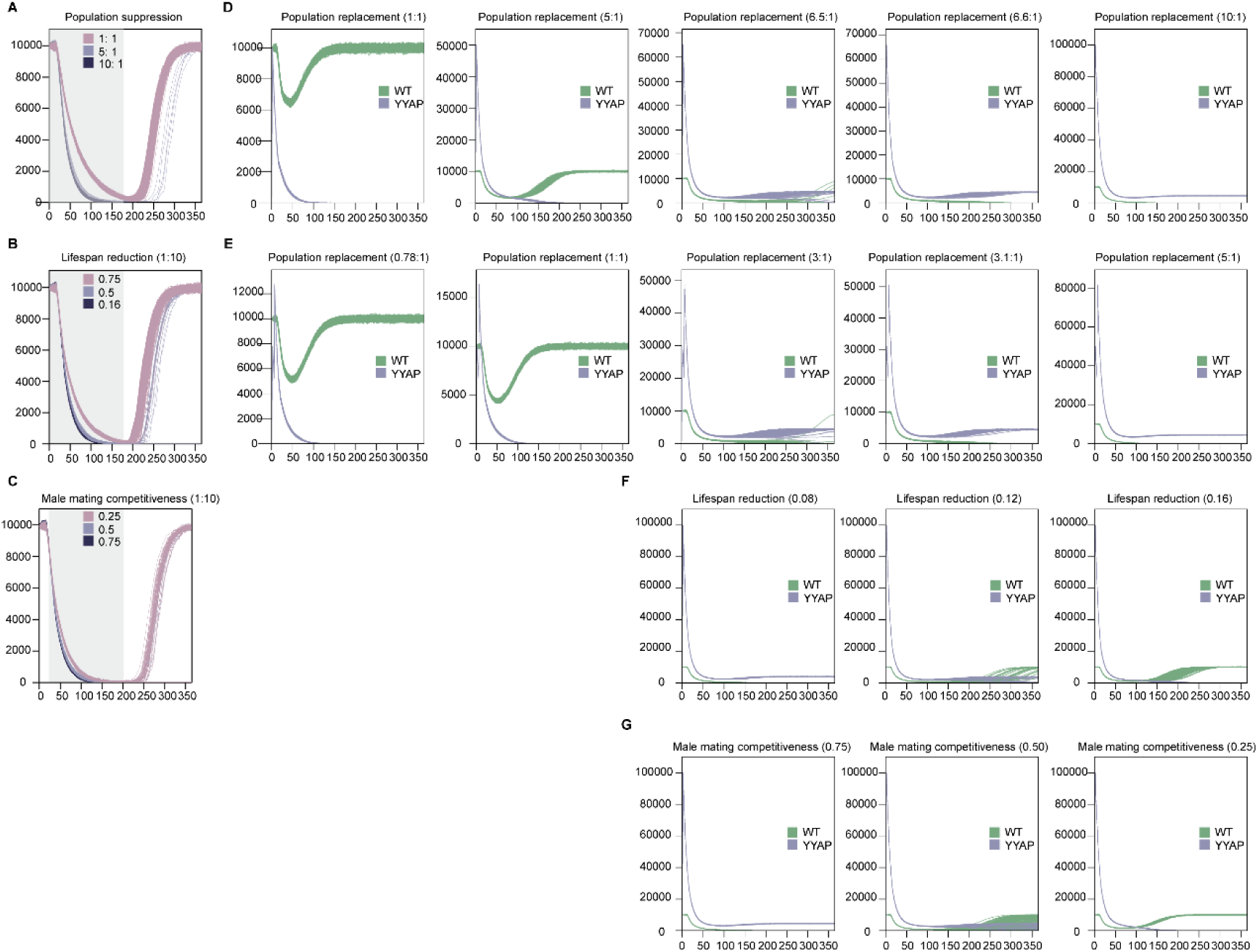
Model-predicted effects of YYAP adult releases on *A. aegypti* population density. (A) Population suppression simulations under ideal fitness conditions. Weekly releases were performed over a 6-month period at release ratios (relative to wild type) indicated in the legend. (B and C) Sensitivity analysis of the YYAP system at fixed release rates. (B) At a release rate of 10 YYAP males per wild-type adult per week, suppression remained achievable with a 16% reduction in lifespan. As lifespan reduction increased, suppression failures occurred more frequently. (C) Under the same release rate, 75% male mating competitiveness reliably achieved elimination, whereas 50% competitiveness led to 8 suppression failures out of 1,000 simulations. (D and E) Modeling of population replacement strategies. (D) Under ideal fitness, a single-release strategy was simulated at varying ratios (relative to wild type, as shown in the key). Replacement was not achieved at release ratios <5, but stable replacement occurred at a ratio of 6.6. (E) Under the same conditions, two-release simulations achieved reliable replacement at ratios of 3.1 per release. (F and G) Sensitivity analysis of the replacement strategy at a release ratio of 10. (F) Due to model constraints, simulations were performed with simultaneous changes in male and female lifespan. Replacement was achieved at 92% lifespan (calculated as the mean of male longevity reduced to 84% with female longevity unchanged). Below 88%, some simulations failed to achieve replacement, and no replacement occurred at 84% or lower. (G) When male competitiveness was reduced to 0.5, partial replacement failures were observed. All simulations were performed in a randomly mixed population of 10,000 adult female Aedes aegypti, with 1,000 stochastic realizations per parameter set using the MGDrivE framework (Sánchez et al., 2019).

**Supplementary Table 1.**
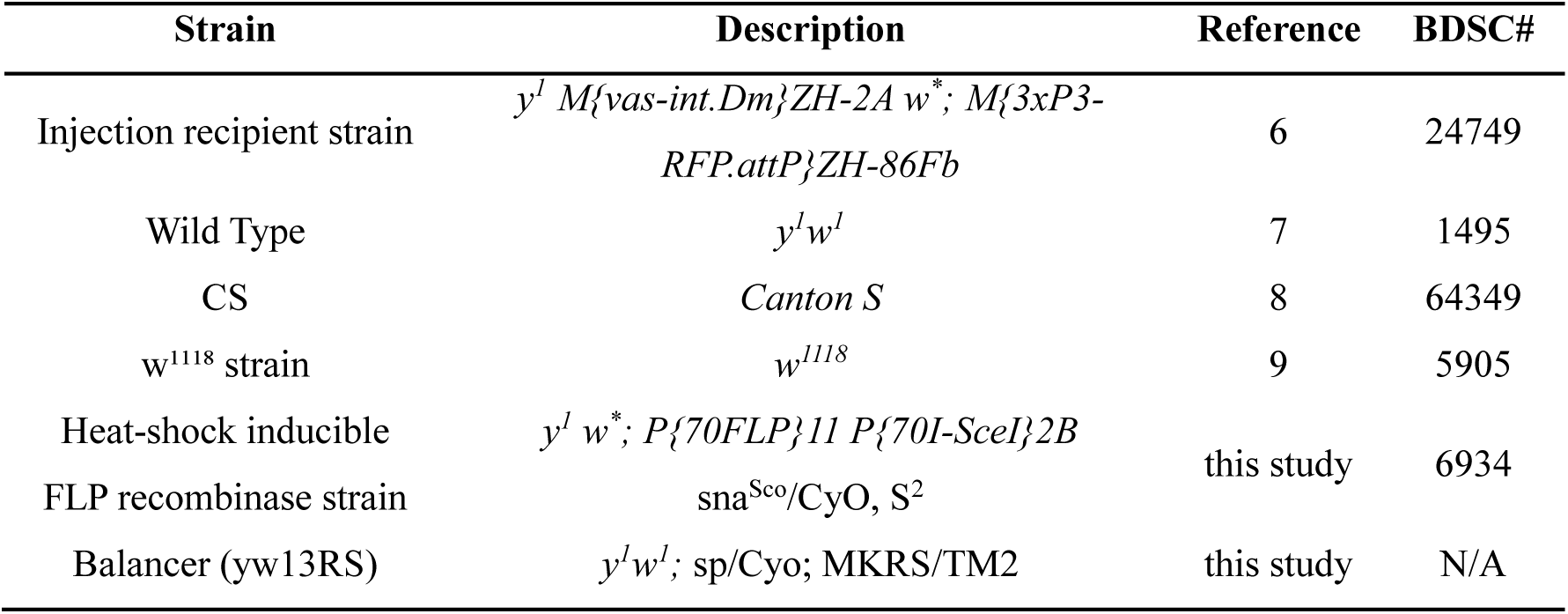
*D. melanogaster* strains used in this study.

**Supplementary Table 2.**
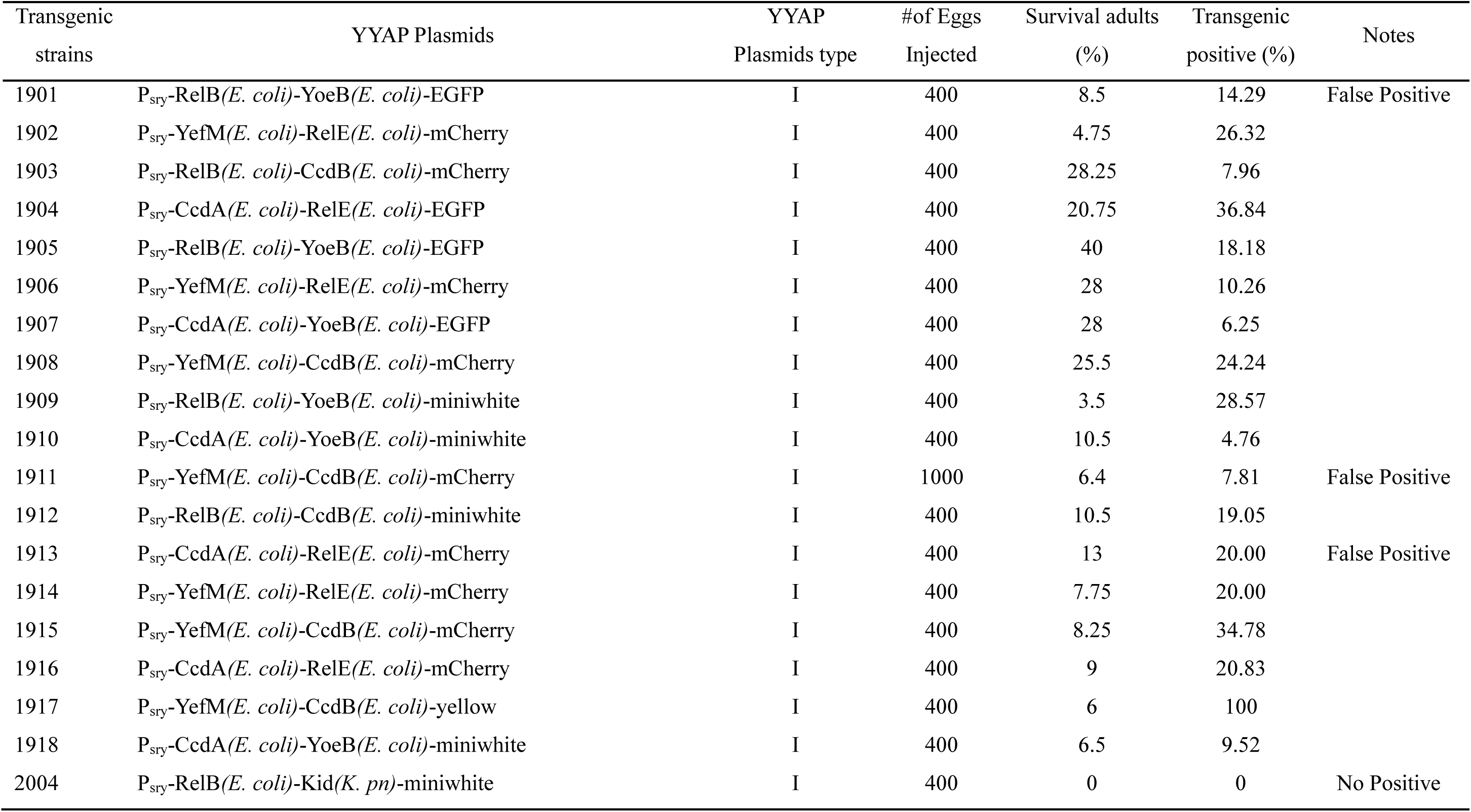
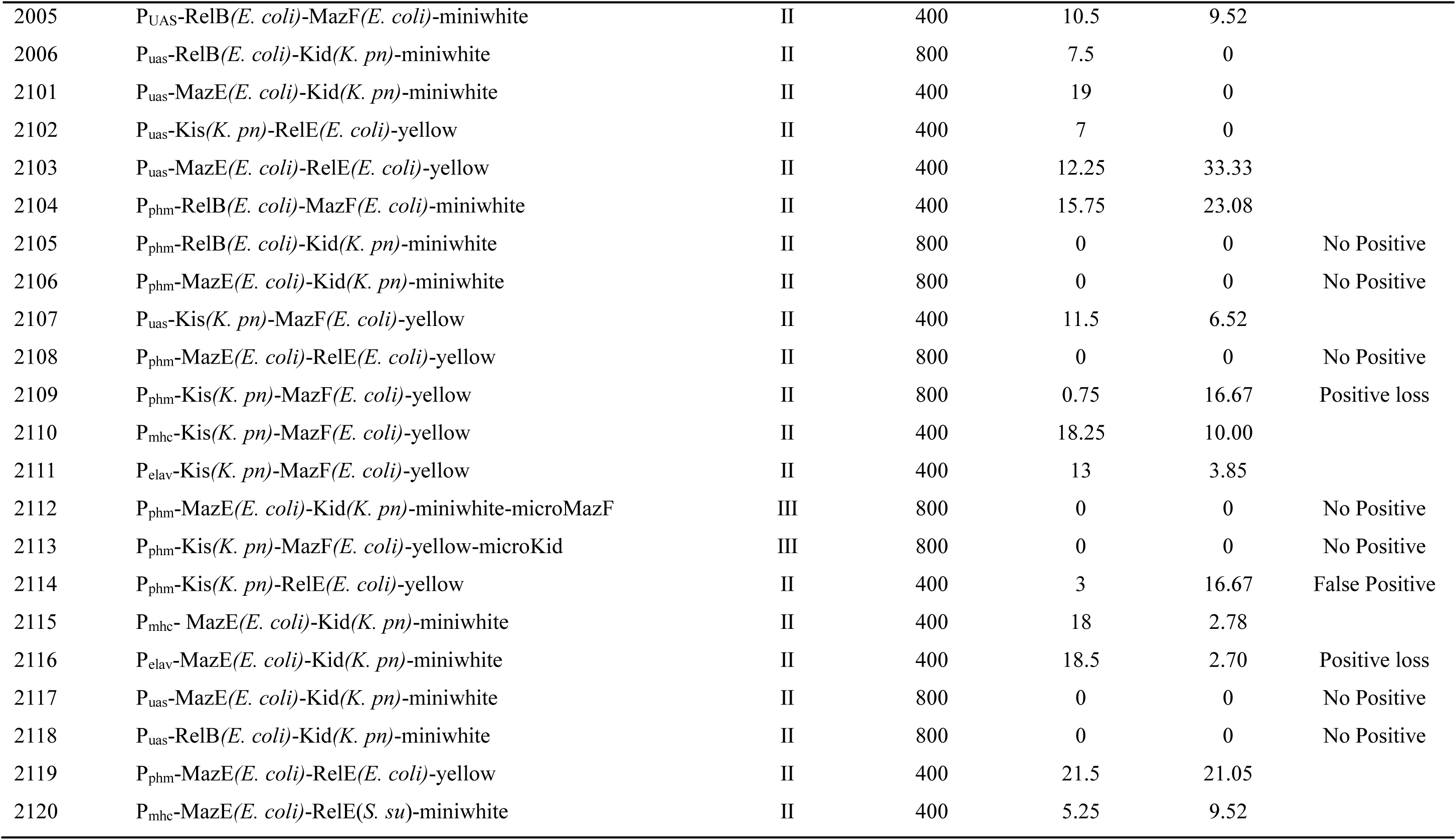
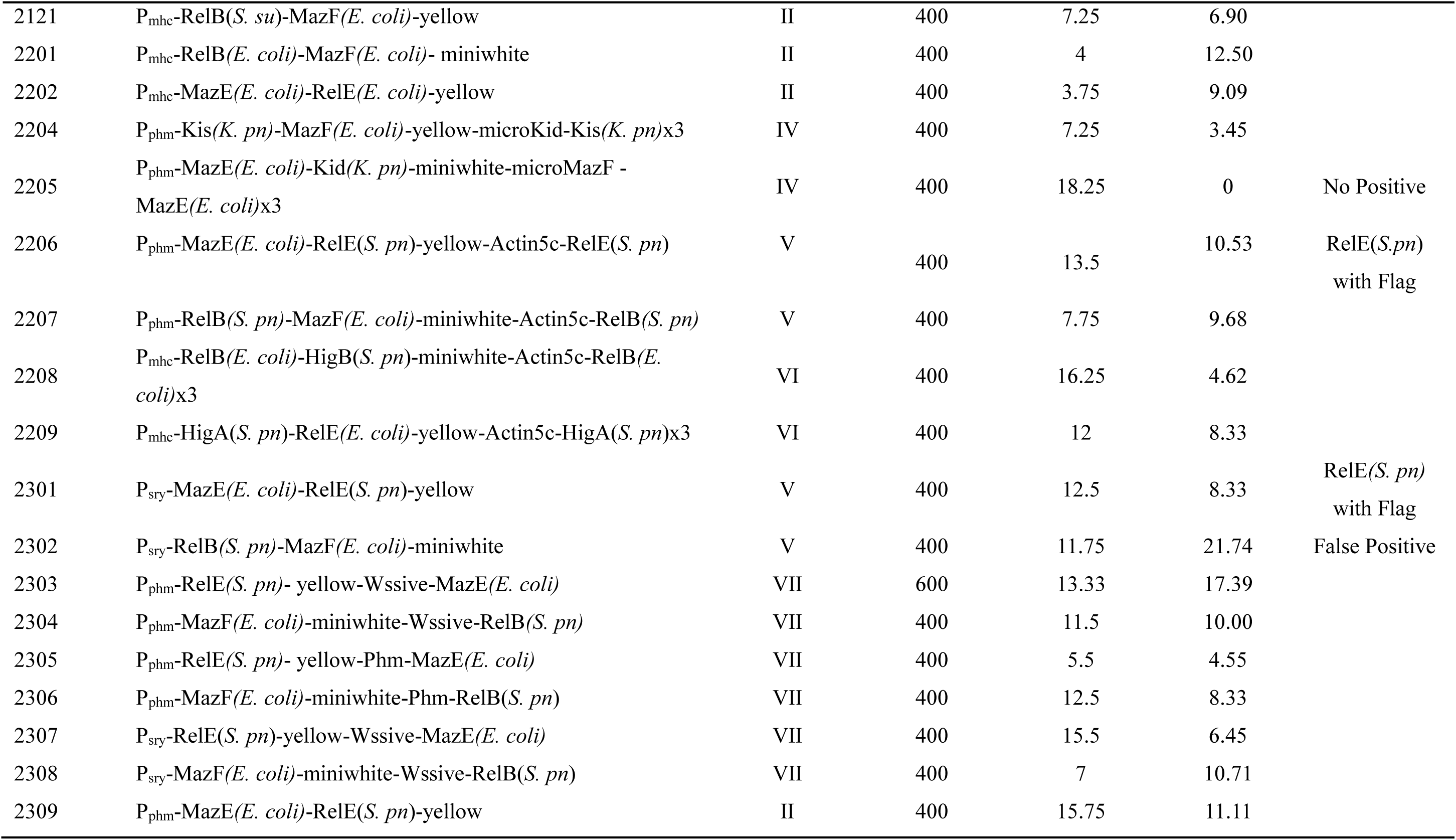
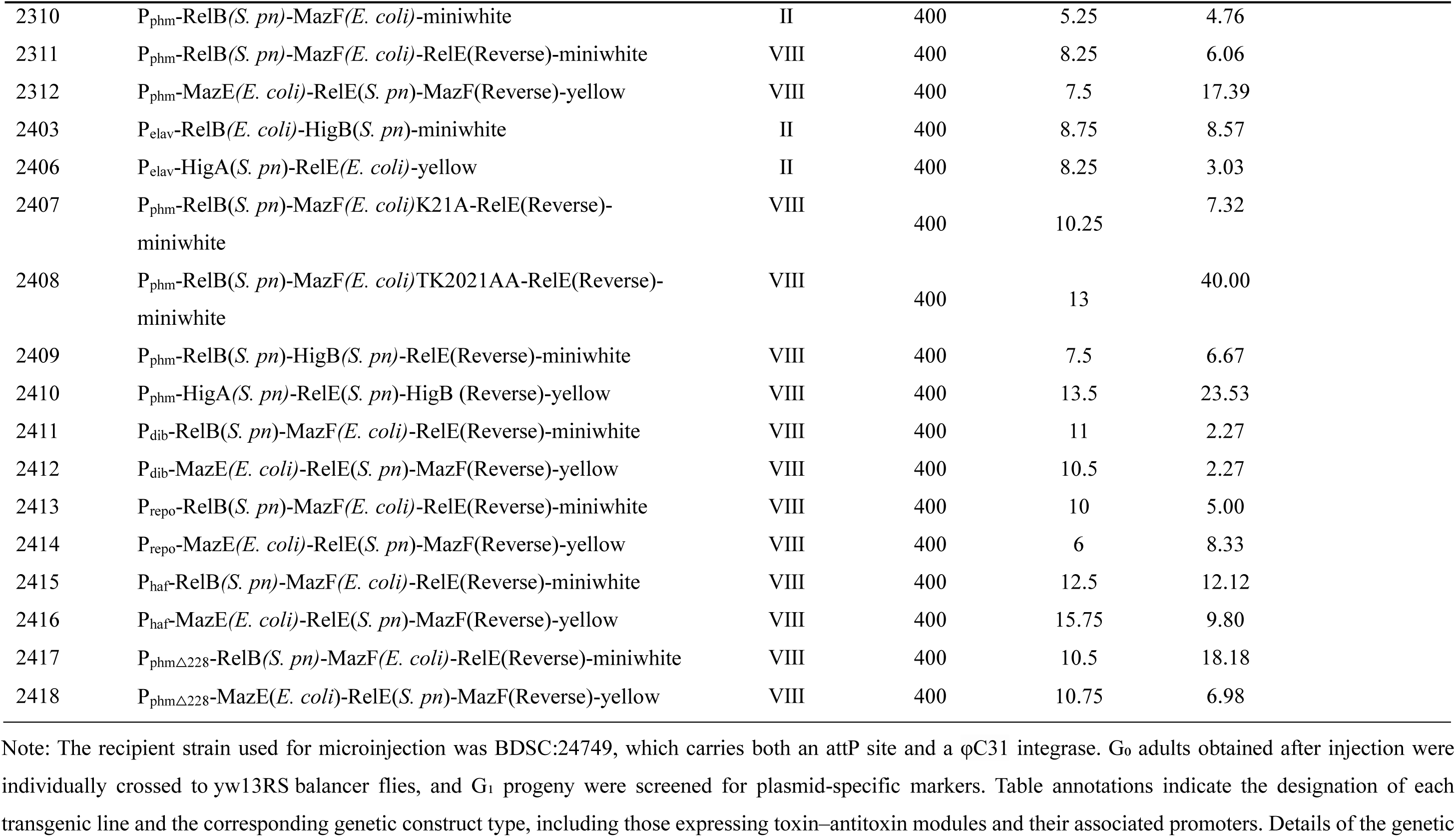
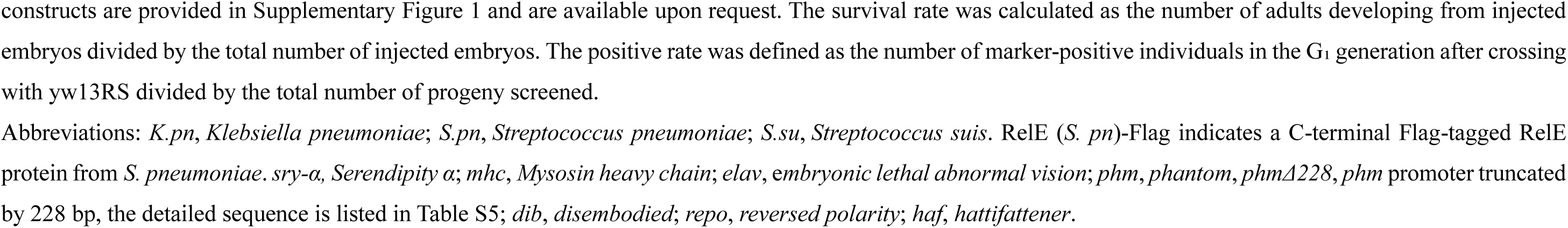
Germline transformation efficiency of YYAP constructs in *Drosophila*.

**Supplementary Table 3.**
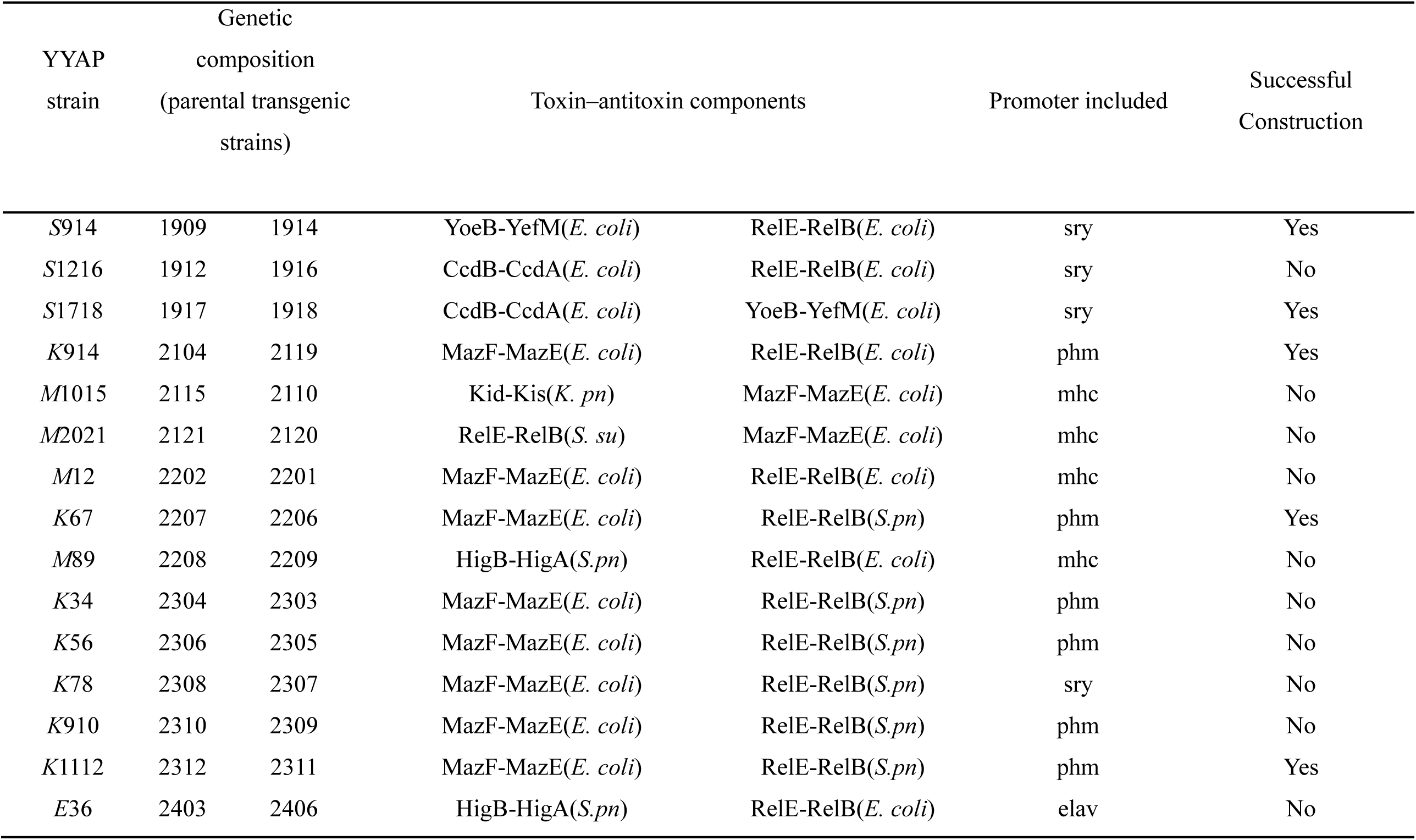
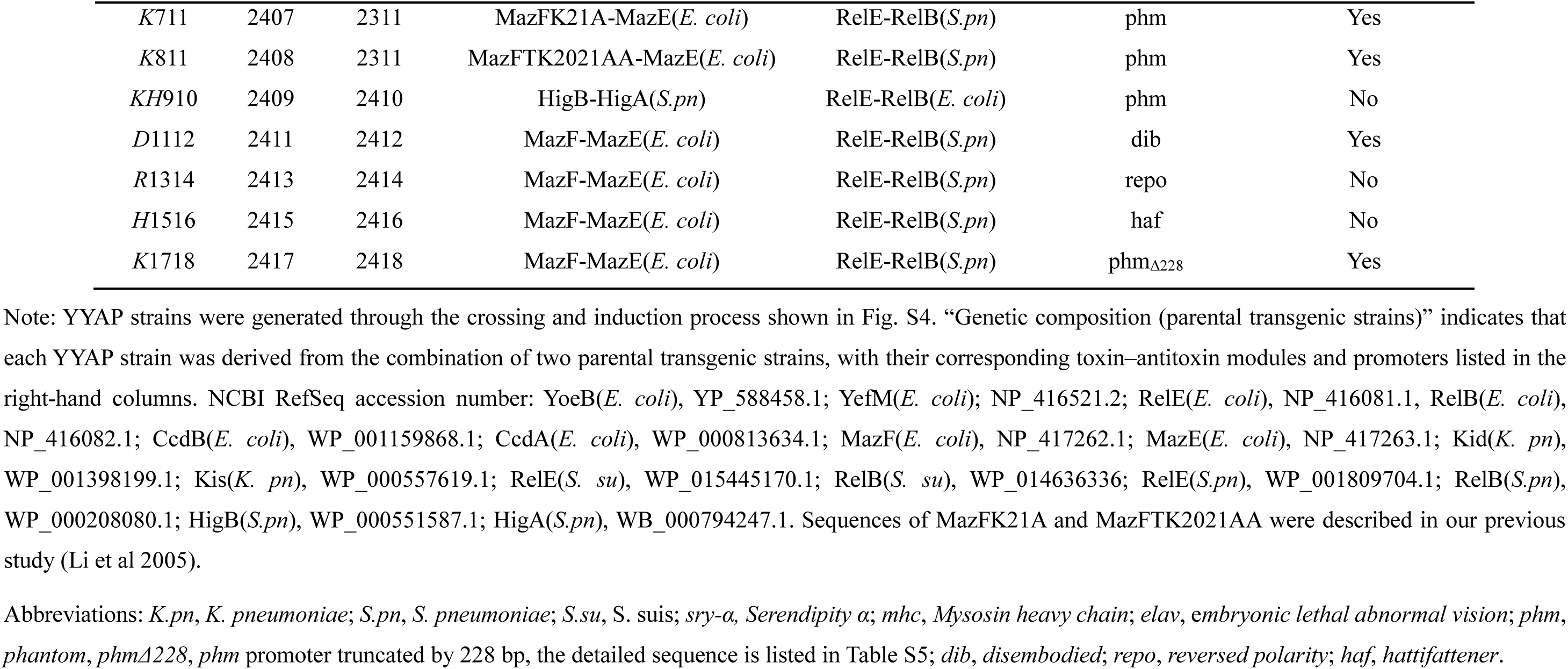
Parental origins and main components of YYAP strains.

**Supplementary Table 4.**
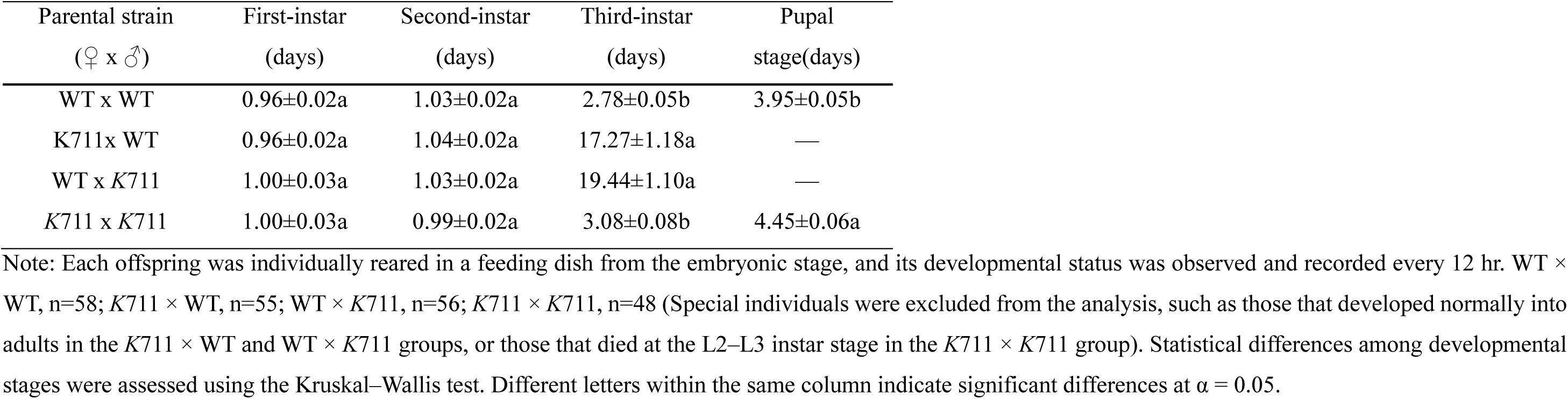
Developmental timing of offspring from different parental crosses.

**Supplementary Table 5.**
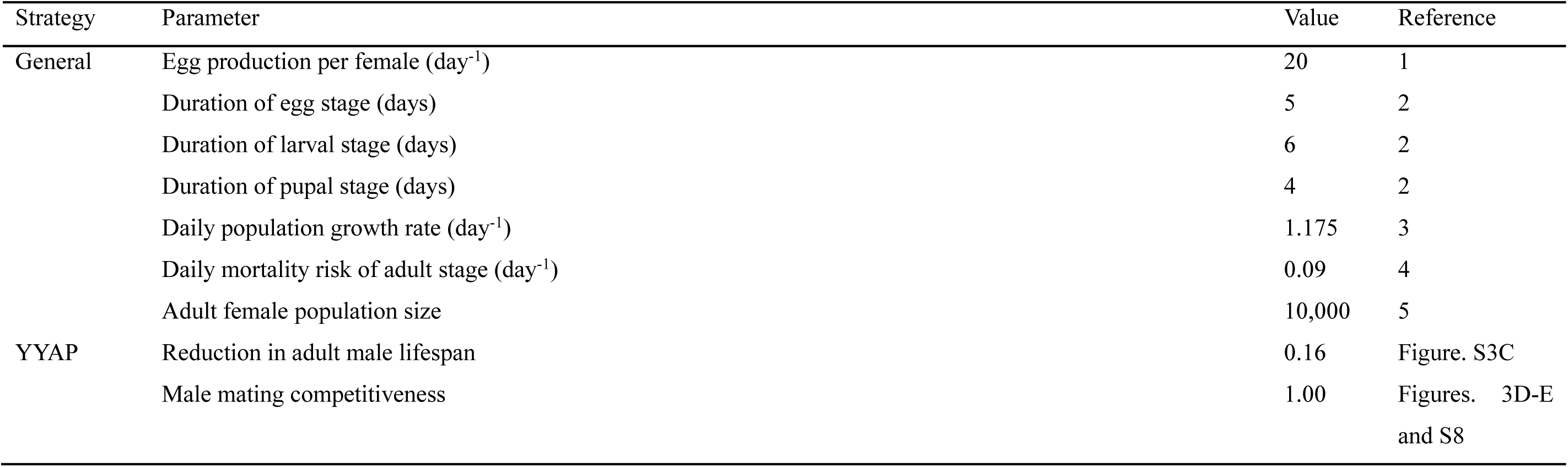
Parameters used in *Aedes aegypti* population suppression and population replacement model.

**Supplementary Table 6.**
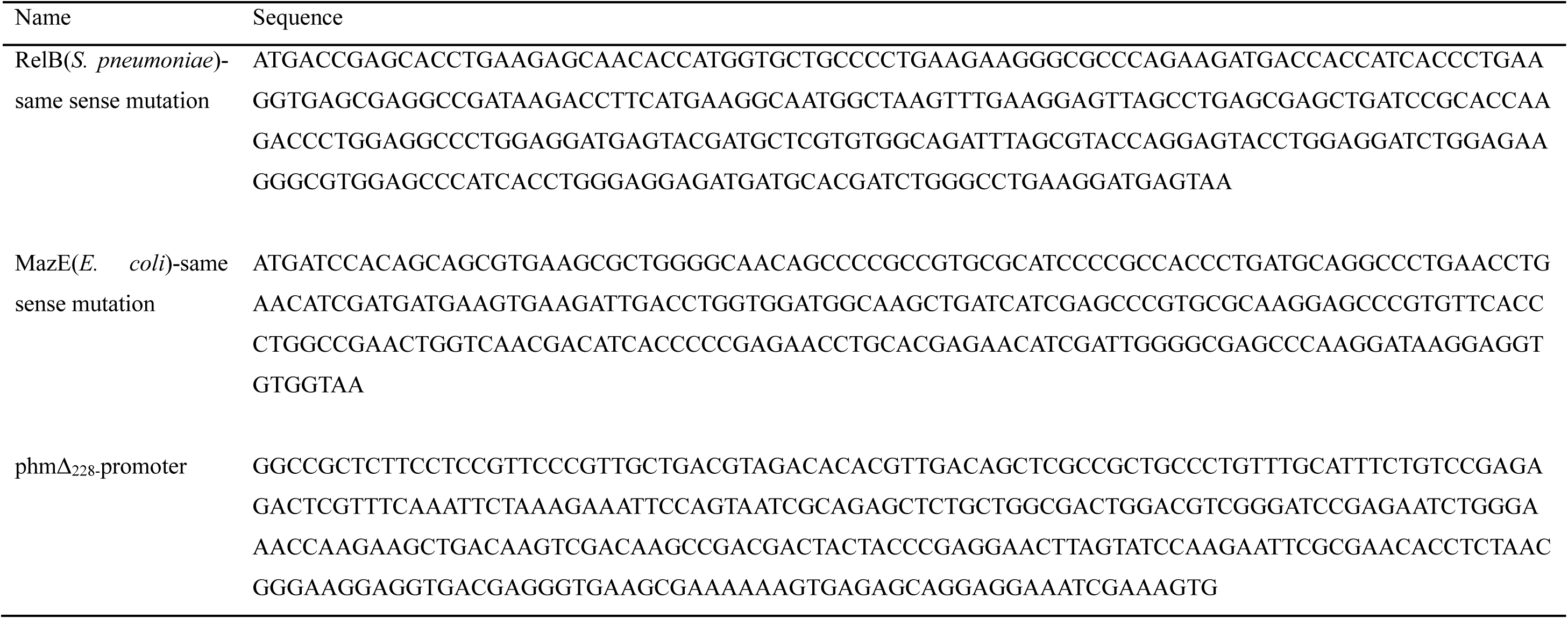
Mutant sequence in this study.

**Supplementary Table 7.**
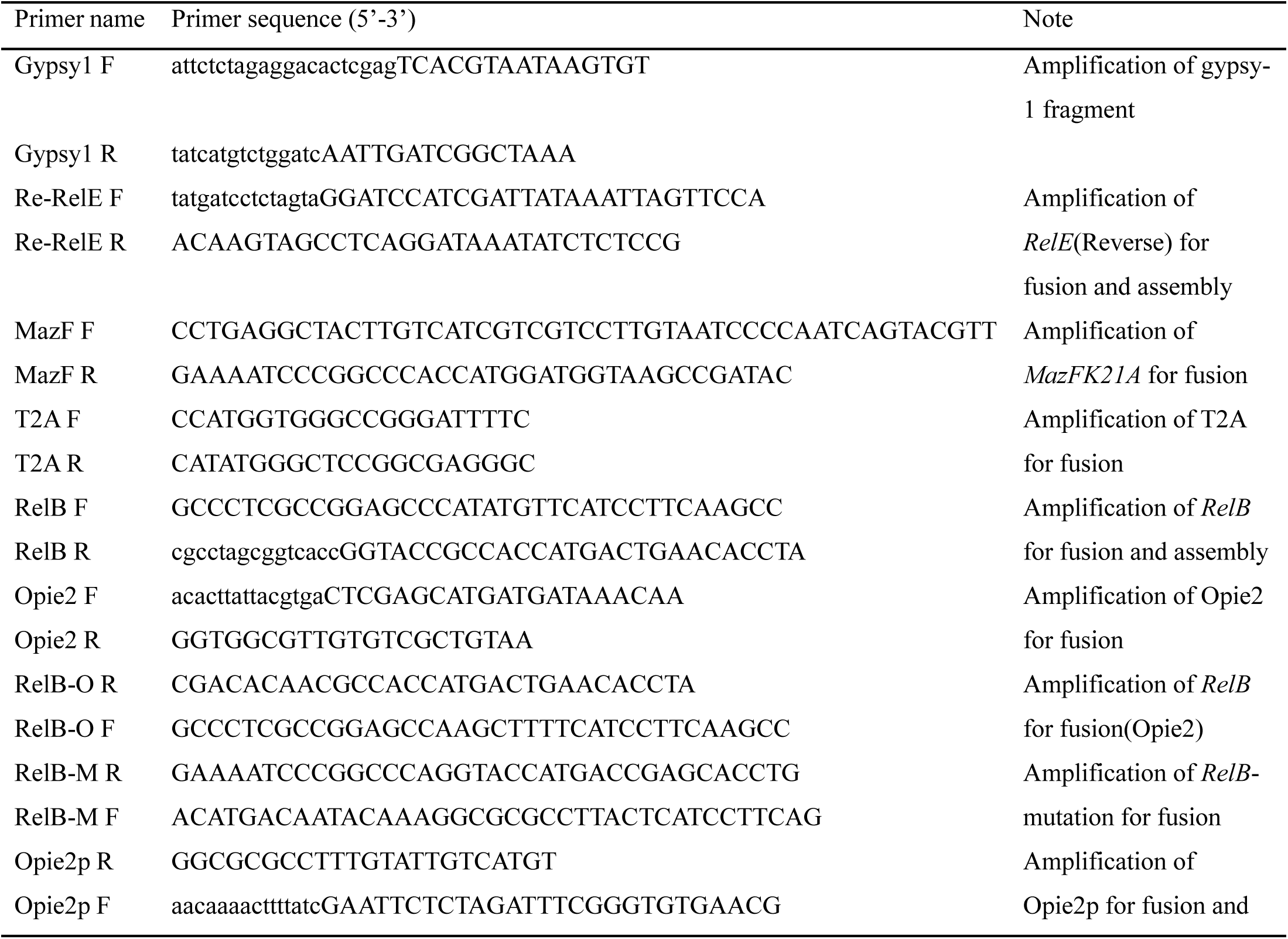
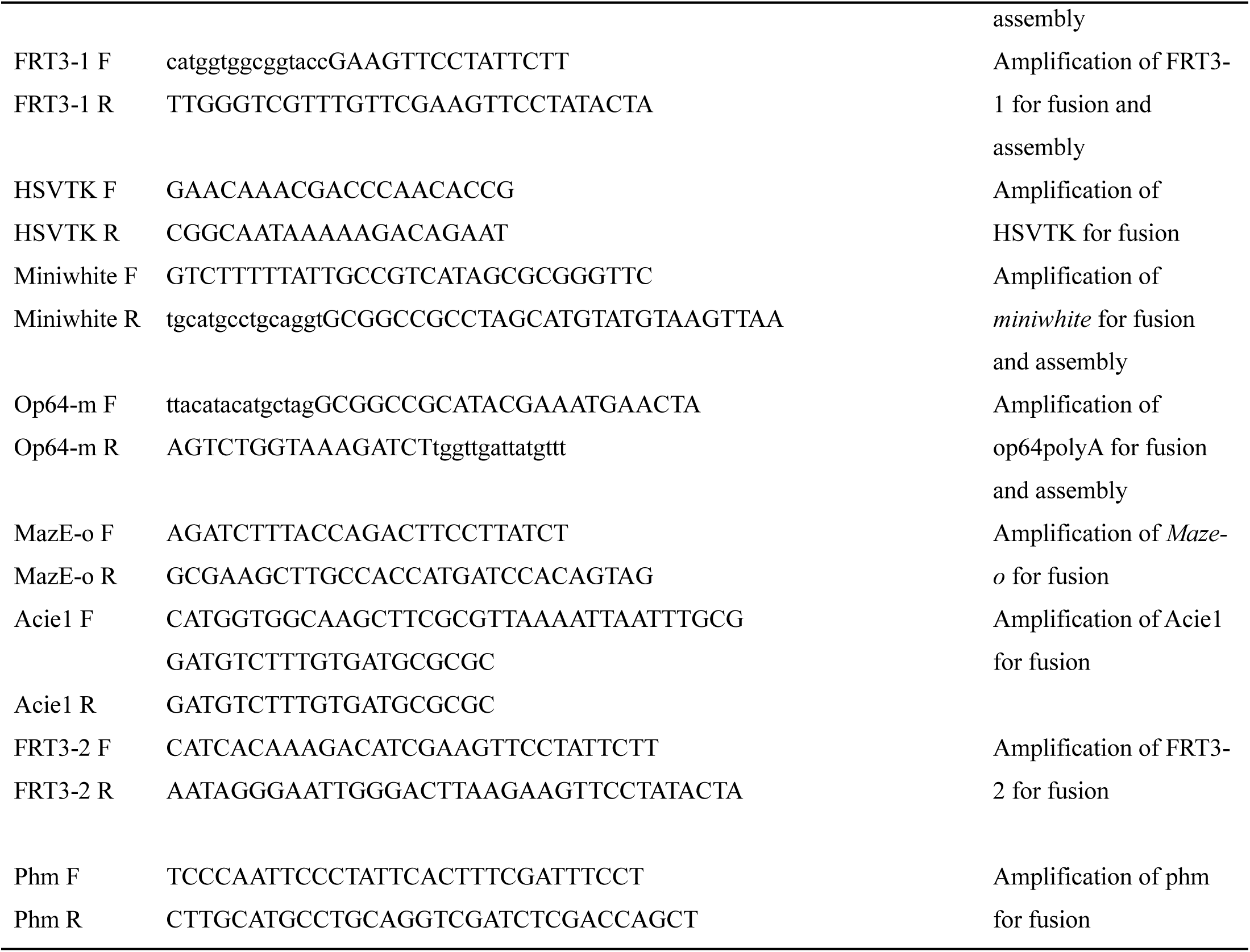
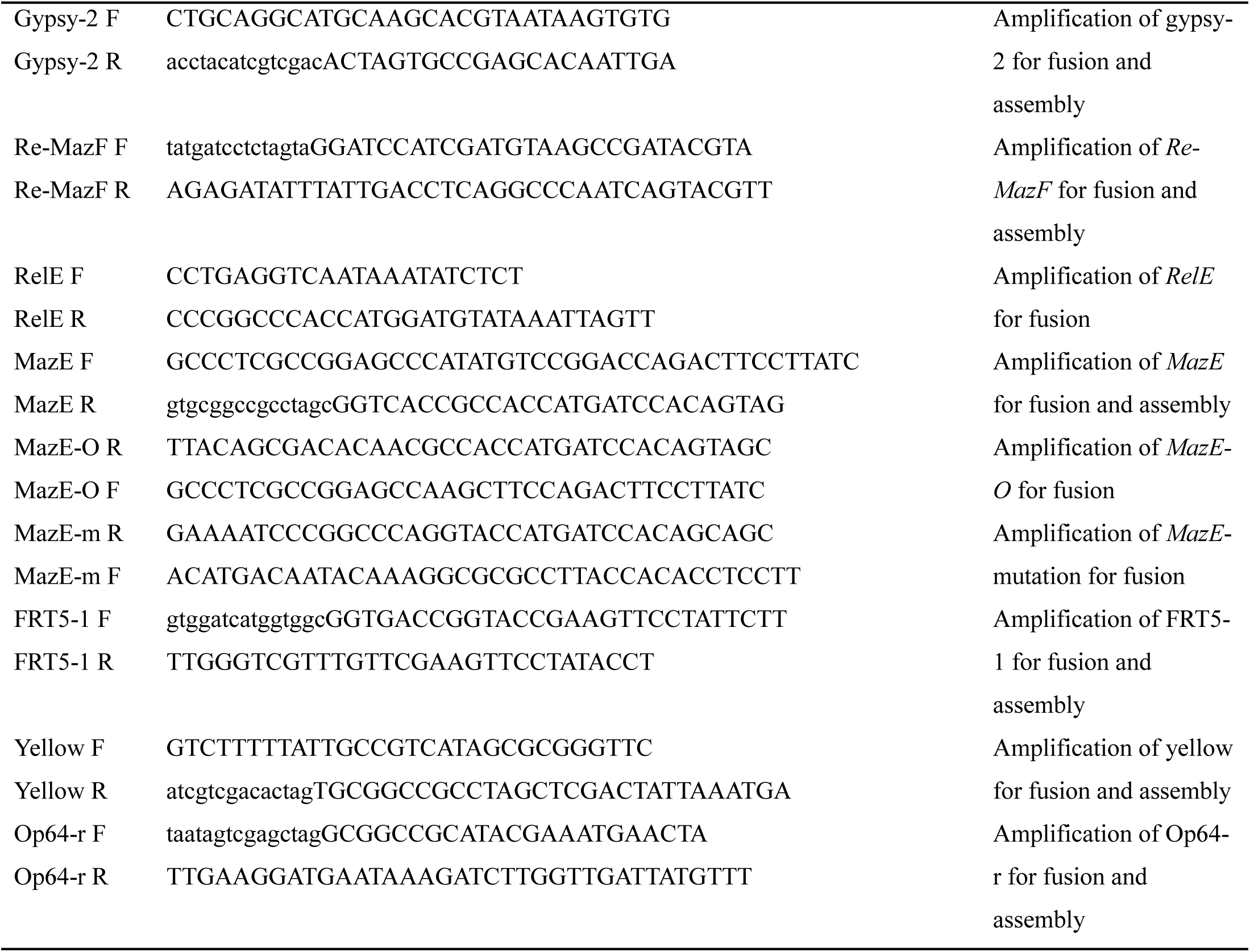
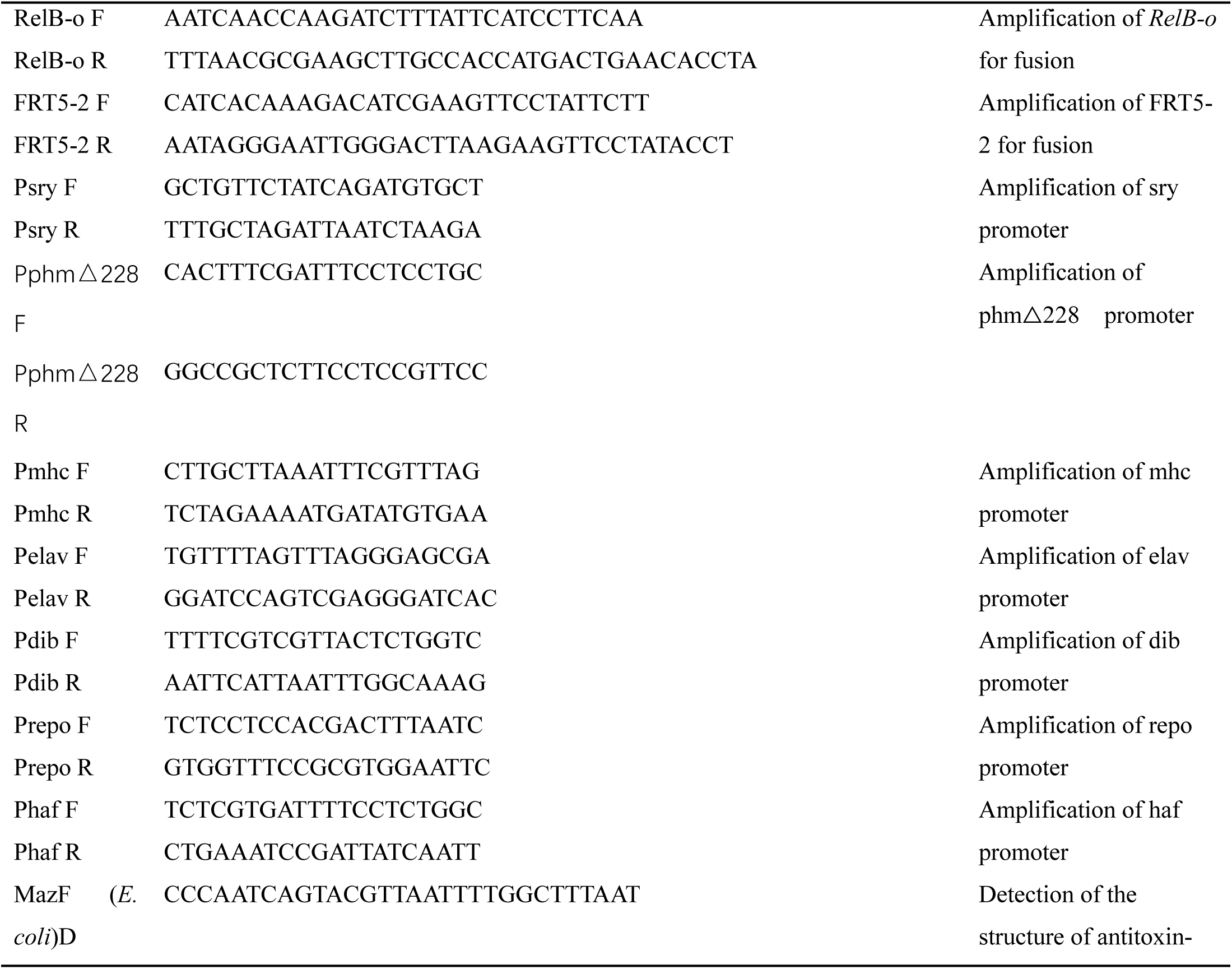
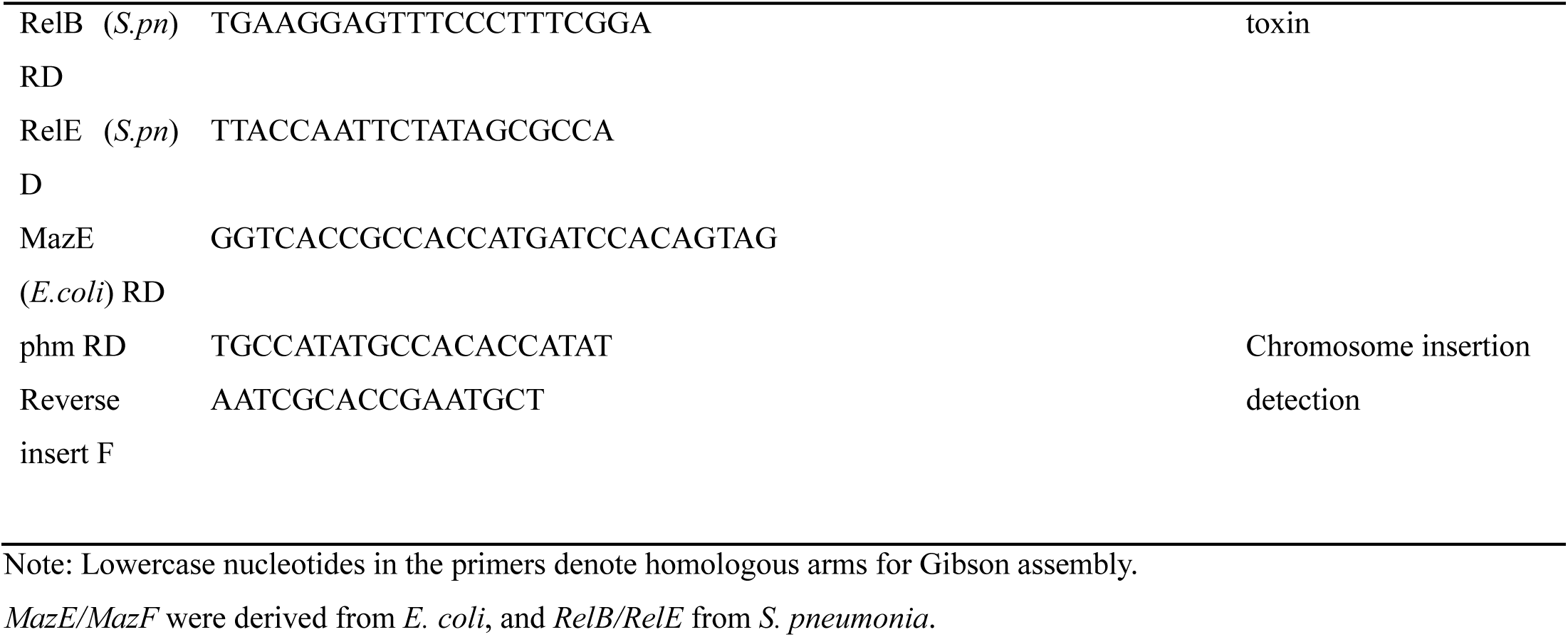
Primer sequences used in this study.

